# 6S RNA tunes virulence, surface immunogenicity, and phenotypic heterogeneity in tissue-invading *Yersinia*

**DOI:** 10.1101/2025.10.07.681022

**Authors:** Oytun Sarigöz, Valerie Diane Valeriano, Ummehan Avican, Sena Gizem Suer, Tugrul Doruk, Sebastian Bauer, Anna Fahlgren, Juliette Griffié, Narmin Hasanzada, Firoj Mahmood, Kristina Nilsson, Angeliki Ntokaki, Helen Wang, Maria Fällman, Kemal Avican

## Abstract

Enteric pathogens must reprogram transcription as they leave the gut lumen and enter tissue, yet this reprogramming has been studied almost entirely in *in vitro* culture systems. Using *Yersinia pseudotuberculosis* mouse infection for *in vivo* RNA-seq, we recovered near-complete transcriptomes from bacteria in the mouse cecal lumen and the associated lymphoid tissue they invade. Tissue-localized bacteria induced the type III secretion system, raised virulence plasmid copy number, and shifted transcription toward alternative sigma factor regulons. The most strongly induced transcript was 6S RNA, among the most conserved noncoding regulators, which sequesters RpoD-containing RNA polymerase but has so far lacked an infection phenotype. Our study shows that 6S RNA acts as a tuner, not a switch: its absence causes T3SS overexpression, increased display of TLR-activating envelope components elevating host expression of proinflammatory mediators, reduced phenotypic heterogeneity in tissue, and delayed colonization. T3SS and 6S RNA transcripts are concentrated in bacteria facing infiltrating neutrophils, defining a local program that couples virulence timing to immune evasion.

**Significance Statement:** Bacteria that cause intestinal disease must switch on virulence systems as they cross from the gut into tissue, yet doing so is costly and increases immune visibility. How pathogens balance this during infection has remained difficult to observe. By resolving near-complete bacterial transcriptomes and spatial organization within infected mouse intestine, we found that the most strongly induced gene encodes 6S RNA, a universally conserved noncoding regulator. Rather than switching virulence on, 6S RNA restrains it: bacteria lacking it overexpress their secretion system, display more immunostimulatory surface antigens, provoke heightened inflammation, and colonize tissue more slowly. For an invading pathogen, more virulence is not always better.

## INTRODUCTION

Diverse enteropathogenic bacteria have evolved sophisticated mechanisms enabling colonization in the gastrointestinal tract of mammalian hosts, expansion to deeper tissue and dissemination to different organs. First layer of the host defence against invading bacteria is through protective function of the intestinal epithelium with associated mucus layer, which serves as a physical barrier to restrict interactions between the lumen and tissue(1). Besides a physical barrier function, intestinal cells also produce and secrete antimicrobial peptides preventing colonization of pathogenic microorganisms and direct contact of microbiota(2–5). Thus, for enteropathogenic bacteria to effectively infect the gastrointestinal tract and pass the epithelial barrier, they must be able to resist antimicrobials secreted by epithelial cells, attacking immune cells and compete with microbiota for available nutrients and space. Once enteropathogenic bacteria have escaped the competitive luminal environment and moved into the mucosa, they must contend with the mucosal immune defence, which includes Peyer’s patches and lymphoid follicles whose overlaying epithelium contains specialized microfold cells (M cells). While some enteropathogens such as *E. coli* (EPEC) crosses epithelial barrier trough interaction via epithelial cells(6), some pathogens take the advantage of M cell-mediated mucosal sampling of antigens to the lymphoid follicles to cross epithelial barrier(7, 8).

Many Gram-negative enteropathogens such as *S. flexneri, Y. pseudotuberculosis, E. coli*, and *S.* Typhimurium have capacity to cross epithelial barrier and colonize in deep tissue sites by escaping from phagocytic killing and manipulating host cell signaling. This capacity relies on their type three secretion system (T3SS), which delivers toxic proteins known as effector proteins to the extracellular milieu or directly into interacting host cells(9). However, regulation of bacterial virulence capacity upon passage from the luminal part of the gastrointestinal tract into neighbouring intestinal tissue has not yet been studied. Even though there are several reports showing need of the T3SS to dysfunction the epithelial barrier at the initial phase of infection, spatiotemporal regulation of virulence for a successful colonization is yet unknown. Furthermore, we lack information of *in vivo* environmental cues that trigger T3SS during infection. Induction of T3SS is known to be costly and lead to growth restriction during *in vitro* induction conditions(10). Hence, there are reasons to believe that temporal and maybe spatial distribution of T3SS expression is important for infecting bacteria.

The aim of this study was to investigate regulation of global gene expression during *in vivo* translocation of an enteropathogen from lumen to intestinal tissue utilizing *in vivo* RNA-seq of the food borne pathogen *Y. pseudotuberculosis*. This model pathogen provides an attractive mouse intestinal infection model where it after oral infection colonizes cecal lymphoid follicles(11). This compartment abounds in immune cells, but *Y. pseudotuberculosis* can resist this defence via its plasmid-encoded T3SS effector proteins, *Yersinia* outer proteins (Yops) that are injected directly into interacting phagocytes(12). Our previous *in vivo* gene expression analysis of mouse cecal tissue biopsies from early and chronic phases of infection by this pathogen revealed transcriptional reprogramming from virulent to persistent mode. This involved a high expression of T3SS during acute infection followed by significant downregulation of T3SS virulence genes, together with an upregulation of genes encoding proteins important for adaption to new environments during the chronic phase of infection(13). In this study, we resolve *in vivo* transcriptional regulation of *Y. pseudotuberculosis* spatially by comparing bacteria in the caecal lumen with those that have colonised the adjacent lymphoid compartment. Here we found that bacteria in the two compartments display distinct transcriptional programmes, with virulence genes and virulence plasmid copy number elevated in tissue where oxygen availability acts as one environmental cue. Further, genes induced on tissue entry largely lacked RpoD promoter motifs and were enriched for stress sigma factor regulons, accompanied by pronounced induction of the non-coding 6S RNA. We show that 6S RNA tunes this programme: its deletion caused T3SS overexpression, increased display of immunostimulatory envelope components, an elevated host inflammatory response, and delayed tissue invasion. Spatial gene expression analysis of infected tissue sections revealed that this programme is restricted within the lymphoid foci, with T3SS and 6S RNA expression concentrated in bacteria adjacent to infiltrating immune cells. Beyond this transcriptional programme, single-colony analysis of bacteria recovered from infection showed that 6S RNA also shapes the diversity of physiological states within the population.

## RESULTS

### Improved bacterial RNA enrichment enables full *in vivo* transcriptome coverage of spatially distinct *Y. pseudotuberculosis* in the cecum organ

To understand transcriptional programs enabling *Y. pseudotuberculosis* to translocate from the intestinal lumen into cecal tissue, we aimed to perform *in vivo* transcriptomics of infecting bacteria localized in tissue and lumen. For this, we orally infected FVB/n mice with low dose of bioluminescent *Y. pseudotuberculosis* YPIII. The infection was monitored with *In vivo* imaging system (IVIS). At two days post-infection (dpi), mice were sacrificed and isolated cecums were subjected to dissection to separate bacteria-containing deep tissue (lymphoid follicles) and associated luminal interphase biopsies (**Fig. 1A**). The separated cecal biopsies from the two compartments were individually homogenized in RNAprotect to stabilize RNA for subsequent determination of *in vivo* transcriptomes of *Y. pseudotuberculosis*. Notably, to ensure sampling of a “well-isolated tissue sample” not containing any luminal content traces, we dissected lymphoid follicle distant from luminal interface. Hence, samples from luminal interface, will therefore likely include some tissue bacteria associated with the interface lining. The two types of compartments will from here be designated ‘tissu’ (cecal lymphoid tissue) and ‘lumen’ (cecal luminal interface).

**Fig. 1.**
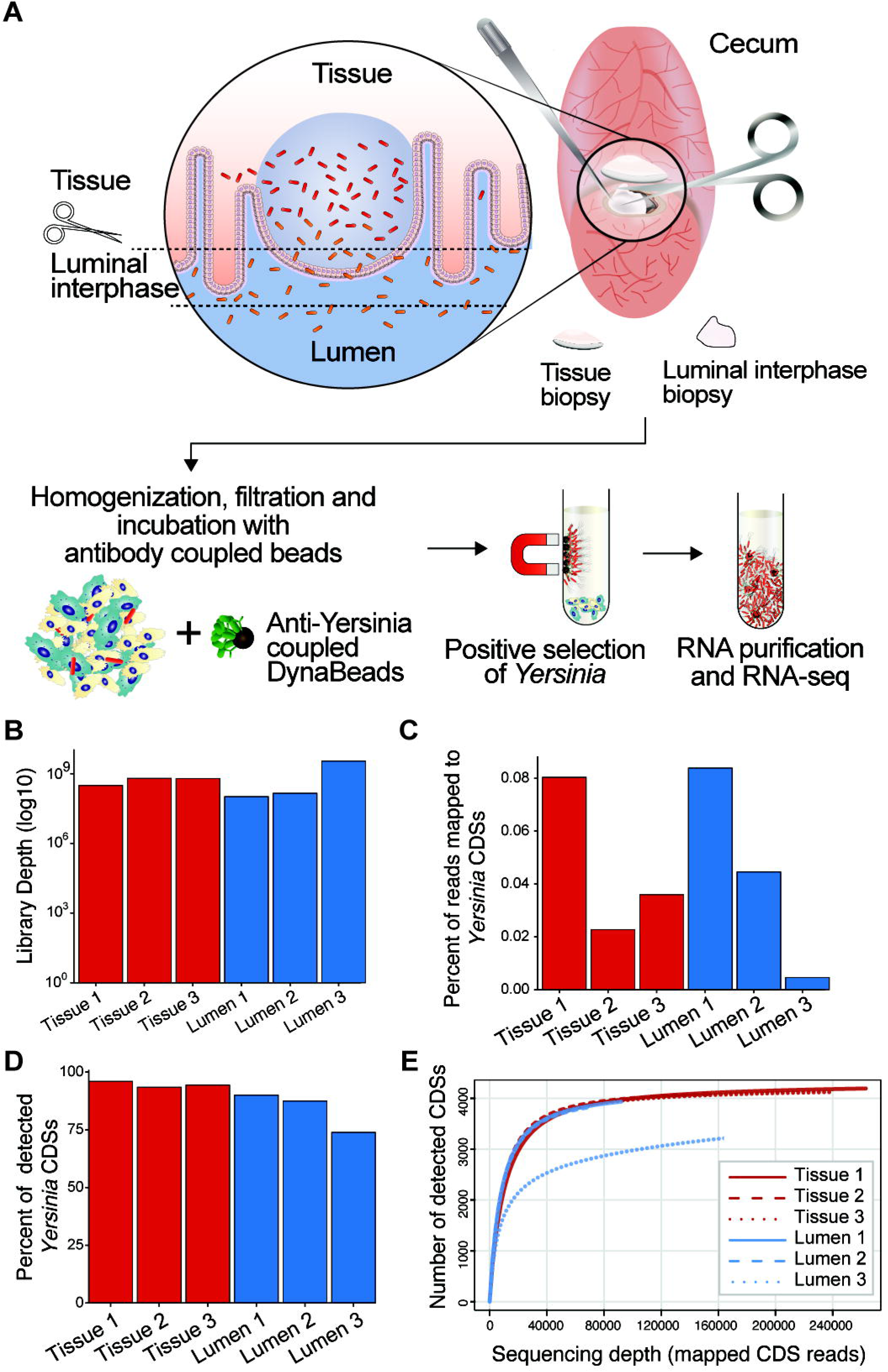
Method for retrieval of high coverage transcriptome profiles of *Y. pseudotuberculosis* in intestinal lymphoid tissue and associated lumen. A Schematic illustration of the procedure with separation of tissue and luminal compartments of mouse cecal lymphoid follicles. Cecums were isolated from mice infected with light emitting *Y. pseudotuberculosis* YpIII/pIBX (expressing the *luxABCDE* operon), allowing localization of the site of infection in the organ. At 2 dpi, cecums were isolated and cecal lymphoid tissue and the associated luminal interface were separated using a scissor whereafter resulting specimens were placed in RNA protect. After homogenization and size filtration, bacterial cells were enriched with anti-*Yersinia* coupled Dynabeads and thereafter subjected to RNA isolation, library preparation and RNA-sequencing. B Total sequencing library depths generated for lymphoid tissue (red) and lumen (blue) samples. C Percentage of RNA-seq reads mapped to *Y. pseudotuberculosis* CDSs. D Transcriptome coverage of *Y. pseudotuberculosis* in lymphoid tissue (red) and lumen (blue) samples. The coverage is calculated as percentage of CDSs with at least one read mapped to the coding sequence. E Rarefaction analysis showing sequencing depth and species richness for tissue (red solid and dashed lines) and lumen (blue solid and dashed lines) replicates. Total RNAs extracted from tissue and luminal compartments of 3 mice cecums were used for RNA-seq.

In our previous study where we analysed *Y. pseudotuberculosis in vivo* transcriptomes and compared gene expression during early and persistent infection in mouse cecum, we could not reach full coverage of the transcriptomes(13). Therefore, we here included an additional bacterial cell enrichment step for retrieval of a complete *Y. pseudotuberculosis* transcriptome from the complex samples populated with host cells and microbiota. For this, we employed anti-*Yersinia* antibodies coupled to Dynabeads to positively select bacteria from cecal biopsy homogenates in order to increase the proportion of *Y. pseudotuberculosis* RNA in the samples. The *Yersinia*-enriched fractions were subjected to total RNA extraction followed by depletion of bacterial rRNA and mammalian rRNA and poly(A) RNAs prior to RNA-seq library preparation and sequencing (**Fig. 1A**). For data analyses, sequencing reads from three biological replicates of tissue and luminal samples were aligned to *Y. pseudotuberculosis* genome with strict alignment parameters allowing only *Y. pseudotuberculosis* specific transcripts to be aligned, as described previously(13). Read mappings of up to 3.5 billions reads from each RNA-seq libraries showed that the addition of the bacterial cell enrichment step resulted in average of 0.082% with bacterial mRNA (**Fig. 1A, C**) compared to previous average of 0.002%. This represents up to 41 times enrichment of bacterial mRNA in the samples in comparison to previous work(13). The percentage of coding sequences (CDSs) with at least one mapped read was increased to nearly 100% for both tissue and lumen located *Y. pseudotuberculosis* except lumen replicate 3 (**Fig. 1D**). Despite a library depth of 3.5 billion reads with 6.9 million reads mapped to the *Yersinia* genome, very few reads mapped to *Yersinia* CDSs, with the fewest CDSs detected in lumen replicate 3, most likely reflecting a very low bacterial load in this specimen. This result suggests that our sequencing depth was sufficient to capture overall transcriptomic profiles of tissue and lumen located *Y. pseudotuberculosis.* The efficiency of our sequencing depth is further confirmed with rarefaction analysis, which illustrates clear saturation of detected genes across all samples except lumen replicate 3 (**Fig. 1E**). Hence, the optimized protocol allowed recovery of full coverage *Y. pseudotuberculosis in vivo* transcriptomes, which will benefit further analyses allowing global analyses involving both high and low expressed mRNAs. Further, with respect to the nature of the samples where the transcriptomes are expected to have both similarities and differences, full coverage transcriptomes will increase chances to identify critical players enabling bacteria to exist in the different compartments.

### The cecal tissue environment promotes *Y. pseudotuberculosis* to increase virulence plasmid copy number and T3SS

Given that environments encountered by enteropathogenic bacteria before and after crossing the epithelial barrier differ significantly in terms of structure and inhabitants, we hypothesized that bacterial physiology in these compartments would differ. To validate the consistency between the tissue and lumen samples, we generated density plots of gene expression profiles (**Fig. 2A**). Overlapping distributions confirm that the three replicates of each sample type share similar profiles, with the exception of lumen replicate 3, which had the lowest number of detected CDSs. Furthermore, Spearman’s correlation analysis confirmed significant and high level of correlation between the tissue (R=0.84-0.86) and lumen replicates (R=0.74-0.55), indicating biological consistency across the dataset (**Fig. 2B**). Notably, lumen replicate 3 had the lowest correlation with other two replicates (R=0.549 and 0.555). The principal component analysis (PCA) on the global transcriptomes of *Y. pseudotuberculosis* from 6 biological samples distinctly separated the tissue replicates from the luminal replicates (**Fig. 2C; Supplementary Table 1**). This indicates a successful separation during the dissection of tissue and associated luminal interphase biopsies, and even more important, it shows that upon entering the tissue compartment this enteropathogen establish a physiology distinct from its siblings in lumen located millimetres away. This implies that environmental cues in the tissue compartment forces changes of *Y. pseudotuberculosis* global transcriptome for adaption of a physiological state suited for tissue colonization. Lumen replicate 3 separated markedly from the other two lumen replicates along PC1, which captured the majority of transcriptome-wide variance (67.3%) as shown by the scree plot, identifying it as a clear outlier (**Fig. 2C**).

**Fig. 2.**
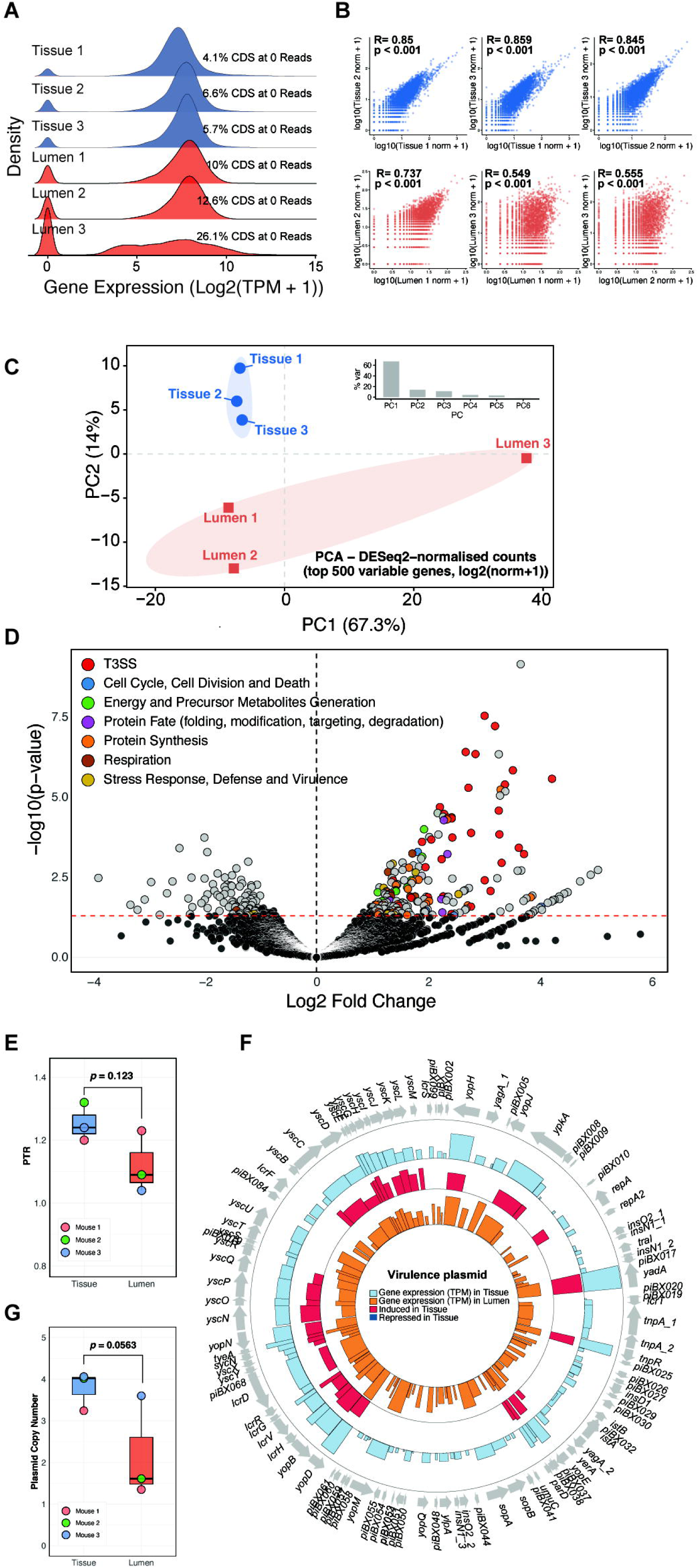
The cecal tissue environment boosts *Y. pseudotuberculosis* proliferation and virulence. *Y. pseudotuberculosis in vivo* gene expression in cecal lymphoid tissue and lumen were analyzed at 2 dpi. **A** Density plot analysis comparing tissue (blue) and lumen (red) replicates. Density distributions were generated using log (DESeq2 normalized counts + 1). **B** Correlation analysis between tissue and lumen replicates. Spearman correlation was calculated using log transformed normalized counts. **C** Principal component analyses (PCA) of the *in vivo* transcriptomes of *Y. pseudotuberculosis* in tissue and lumen. **D** Volcano plot showing differential expression levels of DEGs and functional annotations of genes, obtained from PATRIC Subsystems, with higher expression in tissue located bacteria compared to bacteria in lumen. Black dots indicate genes filtered out by differential expression filter (log2FoldChange >|0.6|, p-value <0.05). **E** Peak-to-through ratio (PTR), measured with digital droplet PCR, as indicative of *Y. pseudotuberculosis* replication level in tissue and lumen. **F** Circos plot of the *Y. pseudotuberculosis* virulence plasmid pIBX showing mean expression level (TPM) for three biological replicates of bacteria located in tissue (blue) and lumen (orange). From outside to inside the circles correspond to: (1) mean expression level of genes in tissue (light blue), (2) genes that are induced (red) or repressed (dark blue) in tissue, and (3) mean expression level of genes in lumen (orange). **G** Copy number of the virulence plasmid pIBX in *Y. pseudotuberculosis* located in tissue and lumen measured with ddPCR. Genes with higher (log2FoldChange > 0.6, p-value <0.05) and lower (log2FoldChange <-0.6, p-value <0.05) expression levels in tissue bacteria compared to luminal bacteria are indicated in red and dark blue, respectively. The data in **E** and **G** are presented as mean values ± SD from 3 sets of specimens (n=3). Two-sample t-tests were employed to measure statistical significance between tissue and lumen located bacteria in **E** and **G**.

To compare *Y. pseudotuberculosis* gene expression between tissue and lumen, we performed differential expression analysis using DESeq2. Lumen replicate 3 was excluded from the differential expression analysis as a clear outlier. Retaining it would have inflated within-group dispersion for the lumen condition, reducing power to detect genuine tissue versus lumen differences. Using thresholds of fold-change > 1.5 and *p* < 0.05, we identified 315 differentially expressed genes (DEGs), with 215 upregulated in tissue (**Fig. 2D; Supplementary Table S2**). PATRIC Subsystems functional annotation (**Supplementary Table 3**) revealed that tissue-upregulated genes were enriched in virulence (T3SS), stress response and defense, respiration, energy generation, protein synthesis and fate, and cell division (**Fig. 2d**). This indicates that upon crossing the epithelial barrier, *Y. pseudotuberculosis* adapts to the stressful tissue environment by increasing virulence, metabolic activity, protein synthesis, and replication.

Since virulence induction and replication reciprocally regulated in *Yersinia in vitro*(10), their overlapping expression in tissue bacteria prompted us to investigate this further. To compare bacterial replication between the two compartments, we used droplet digital PCR (ddPCR) to determine the peak-to-trough ratio (PTR) on DNA extracted from the same biopsy homogenates used for RNA extraction. The PTR method measures the DNA coverage ratio of oriC to the replication terminus, which reflects bacterial replicative status(14, 15). Mean PTR values did not differ significantly between luminal and tissue bacteria, although all tissue replicates tended to have higher values (**Fig. 2E**). Among the 99 genes on the pIBX virulence plasmid, 34 were upregulated in tissue compared to luminal bacteria, including genes encoding the T3SS basal body, export apparatus, translocon, needle, effectors (Yops), and chaperones (**Fig. 2F; Supplementary Tables 1 and 2**). Moreover, upregulation of the plasmid replication initiator RepA and the ParDE type II toxin-antitoxin maintenance system suggested that an increased plasmid copy number (PCN) contributed to elevated T3SS expression. We therefore measured PCN in both compartments using ddPCR on the biopsy homogenate DNA. Although the higher PCN in tissue bacteria did not reach statistical significance (*p* = 0.0563; **Fig. 2G**), it paralleled the significant upregulation of T3SS and plasmid replication/maintenance genes, supporting that cecal lymphoid tissue promotes adaptation toward a virulent phenotype. This aligns with our previous finding of increased virulence plasmid levels during infection(16), while here we spatially resolved these regulatory patterns across bacteria separated by millimeters. Together with increased virulence plasmid replication upon entering tissue, the concomitant increase in virulence gene expression likely promotes colonization by protecting the bacteria against immune elimination.

### Bacterial *in vivo* transcriptomes result from multiple environmental cues, which differs in tissue versus lumen

The *in vivo* transcriptomes obtained from *Y. pseudotuberculosis* in cecal tissue and lumen show unique gene expressions patterns previously not seen in *in vitro* laboratory setting. This discrepancy is obvious when comparing the transcriptomic profiles of tissue and luminal bacteria with those of *in vitro*-cultured bacteria exposed to single infection relevant conditions where two *in vivo* transcriptomes clearly separates from all *in vitro* transcriptomes (**Supplementary Fig. S1**). Hence, during infection multiple environmental cues are expected to influence gene expression patterns of bacteria that thrive for adaption to the new environment.

Cecal tissue and luminal transcriptomes reflect distinct environments, with tissue-upregulated genes pointing to compartment-specific cues. These included iron-limitation genes encoding a TonB-dependent heme/hemoglobin receptor (YPK_3889) and a hemin-degrading protein (YPK_3890), alongside oxidative/redox stress genes encoding a quinone oxidoreductase (YPK_3856), GrpE (YPK_2974), and DnaK (YPK_3594). Upregulation of alternative sigma factors RpoH (YPK_3976) and RpoE (YPK_1182) further indicates transcriptional reprogramming upon tissue entry. Tissue bacteria also highly expressed numerous hypothetical proteins and regulators, including an XRE family regulator (YPK_2310), a putative sigma-54 modulation protein (YPK_0504), and H-NS (YPK_2074), which downregulate T3SS *in vitro*(17) and represses *rpoS* expression(18). Moreover, *iscR* (YPK_1275), which is highly expressed under *in vitro* aerobic conditions(19), were upregulated in tissue, although not statistically significant. Collectively, these findings demonstrate that *Y. pseudotuberculosis* undergoes extensive, compartment-specific transcriptional reprogramming in response to the distinct environmental stresses of cecal tissue.

### Oxygen availability constitute an environmental cue contributing to T3SS induction

Prompted by the marked upregulation of T3SS in tissue, we re-evaluated the functional annotation of DEGs in luminal and tissue-located bacteria to identify the underlying environmental cues. Tissue located bacteria significantly upregulated *atpD*, *atpF*, and *atpG* (F F ATP synthase subunits), *nuoD* and *nuoF* (NADH-quinone oxidoreductase subunits), *trx* (thioredoxin), and *rpoH* (sigma³²), indicating aerobic respiration and reactive oxygen species (ROS) generation (**Fig. 3A**). Although expression of the F F ATP synthase and NADH-quinone oxidoreductase subunit genes was not uniform within their operons, such discrepancies commonly arise from post-transcriptional regulation via transcript stability and RNA-protein interactions(20–22). Furthermore, transcriptome data from our previous study on *in vitro* stress responses(23) showed similar regulation of *nuoD, nuoF, rpoH, trx, atpD, atpF*, and *atpG* during aerobic versus microaerophilic growth (**Fig. 3B**). Together with the higher oxygen levels in intestinal tissue compared to the lumen(24), this regulation in tissue bacteria suggests that oxygen serves as an environmental cue contributing to T3SS expression in this compartment. To test how oxygen availability affects T3SS induction, we exposed Y*. pseudotuberculosis* to 0–5% oxygen at 37°C under virulence-inducing and non-inducing conditions. T3SS expression increased with rising oxygen levels, peaking at 5% (**Fig. 3C**), the concentration found in mouse cecal tissue(24). qPCR confirmed oxygen-dependent induction of the effector genes *yopE* and *yopH* and the translocator gene *yopD* (**Fig. 3D**; **Supplementary Fig. S2B**). However, elevated oxygen alone failed to induce T3SS under non-inducing conditions (**Fig. 3E, F; Supplementary Fig. S2A**), indicating that increased oxygen acts as a contributing cue for T3SS induction upon crossing the epithelial barrier.

**Fig. 3.**
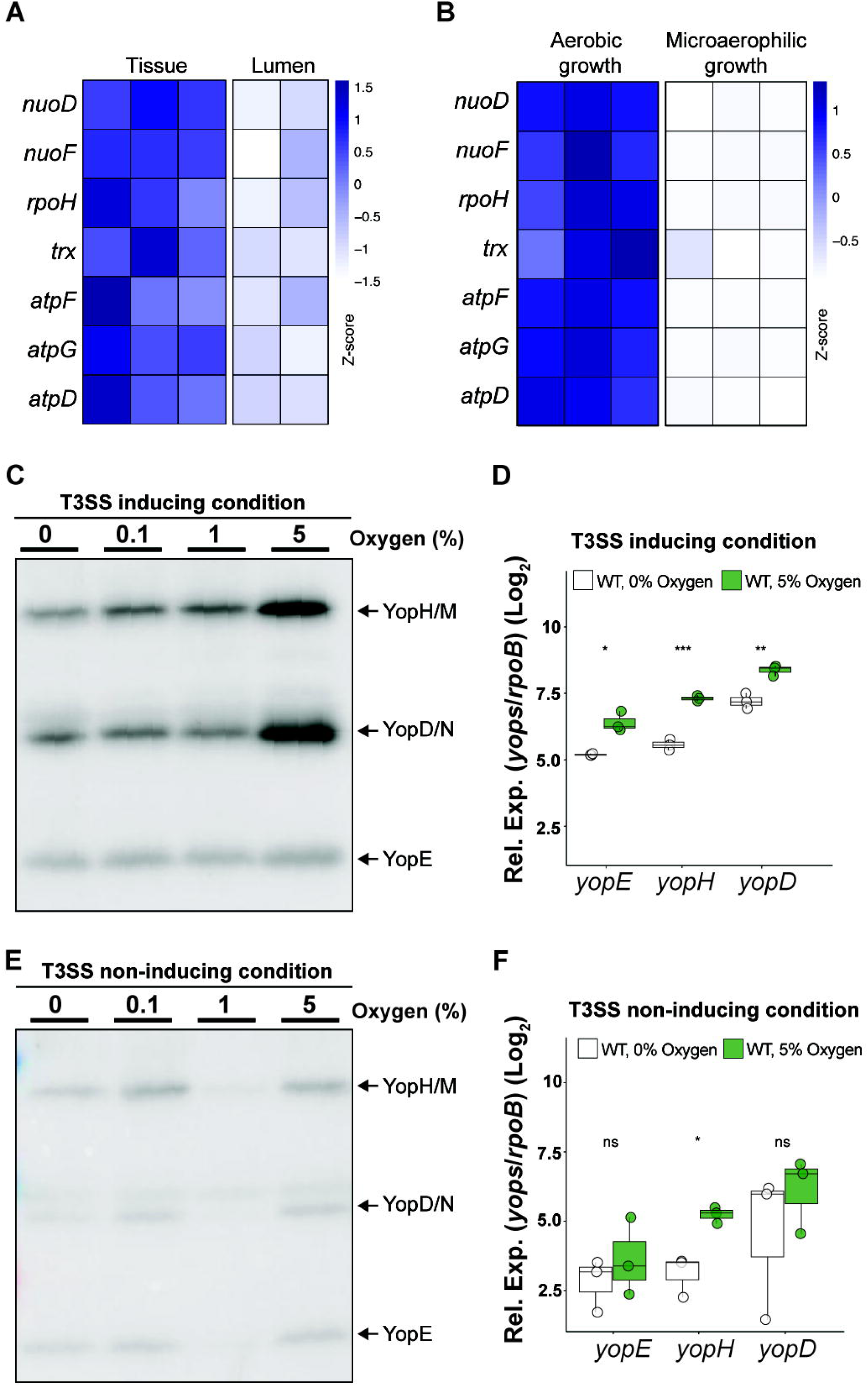
Bacteria in tissue utilize aerobic respiration, a condition required for efficient T3SS induction *in vitro*. **A-B** Z-score heatmaps of aerobic respiration and redox stress genes (*nuoD, nuoF, rpoH, trx, atpF, atpG, atpD*) across the three biological replicates *in vivo* **(A)** and *in vitro* **(B) c** Secretion of T3SS effector proteins in T3SS inducing (37°C, Ca^2+^ depletion) conditions under different oxygen concentrations (0, 0.1, 1, and 5%) for 3 hours. **D** Log Relative expression of *yopE, yopH,* and *yopD* in T3SS inducing condition under 0% and 5% oxygen for 3 hours. **E** Secretion of T3SS effector proteins in non-inducing (37°C) condition under different oxygen concentrations (0, 0.1, 1, and 5%) for 3 hours. **F** Log Relative expression of *yopE, yopH,* and *yopD* in T3SS non-inducing condition under 0% and 5% oxygen for 3 hours. In **D** and **F**, data are presented as log relative expression (Ct values for each gene normalized to Ct values of rpoB). Two sample t tests were performed on log transformed relative expression values. In **D** and **F**, asterisks represent p-value thresholds: *** for p < 0.001, ** for p < 0.01, and * for p < 0.05; ns, non significant. Multiple comparisons and statistical significance were calculated by two-factor ANOVA test (main effects and repeated measures were considered) in Prism 10.

### Tissue-associated bacteria induce 6S RNA and engage the RpoE and RpoH regulons

Besides CDSs, non-coding RNAs (ncRNAs) regulate stress responses, virulence, and biofilm formation by controlling gene expression(25–29). To assess their contribution to transcriptional changes upon tissue entry, we examined well-known annotated *Y. pseudotuberculosis* ncRNAs in luminal and tissue bacteria, alongside *in vitro* logarithmic growth at environmental temperature and T3SS-inducing conditions. DESeq2-normalized read counts revealed significantly higher expression of FMN, CsrC, and 6S RNA in tissue versus luminal bacteria (**Supplementary Fig. S3 and Fig. 4A**). Notably, expression of the global stress regulator 6S RNA (encoded by *ssrS*) was significantly higher in tissue bacteria than during *in vitro* logarithmic growth, but did not differ significantly from *in vitro* virulence-inducing conditions, consistent with 6S RNA induction upon T3SS activation (**Fig. 4A**).

**Fig. 4.**
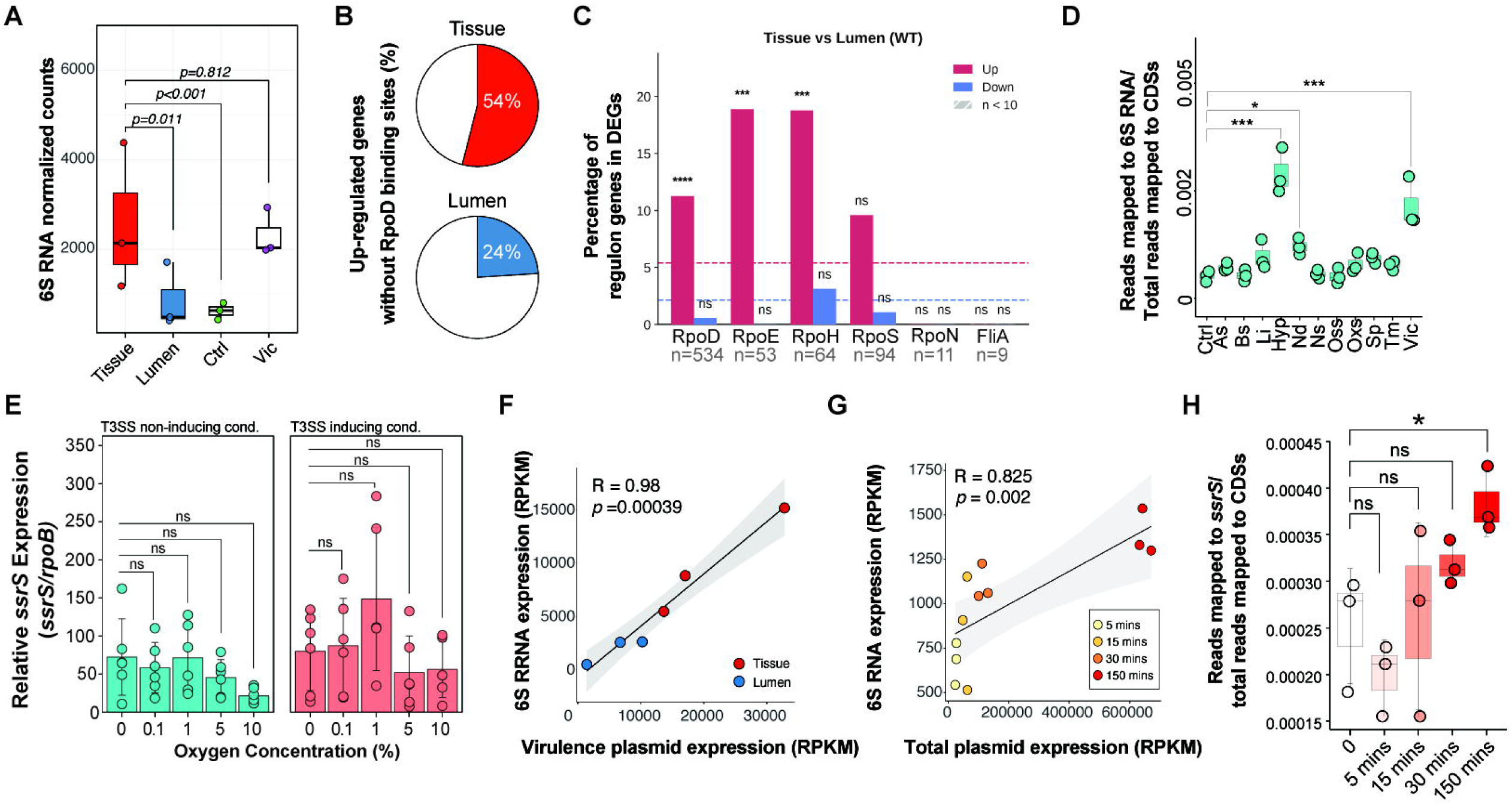
Expression of 6S RNA is increased in tissue-colonized bacteria and linked T3SS. **A** Relative expression of 6S RNA in *Y. pseudotuberculosis* during infection in cecum (lumen and tissue) and *in vitro* logarithmic growth at 26°C (Ctrl) and virulence inducing conditions for 1.5 hour (Vic). Data shown as DESeq2-normalized counts. Data are presented as mean values ± SD from *in vivo* and *in vitro* samples (*n*=3 for all samples except *n*=2 for lumen samples). **B** Percentage of upregulated genes in lumen and tissue located bacteria without a predicted upstream RpoD binding site. **C**Sigma factor–gene interactions were retrieved from RegulonDB (v14.5) and restricted to those annotated as Confirmed or Strong confidence. Interactions were transferred to *Y. pseudotuberculosis* YPIII by reciprocal best-hit sequence alignment. For each regulon, bars show the percentage of member genes significantly up-regulated (red) or down-regulated (blue) in tissue-located relative to lumen-located WT bacteria. Dashed lines indicate the corresponding background fractions of up-regulated (red) and down-regulated (blue) genes across all 3,987 tested genes. The number of YPIII genes assigned to each regulon and present in the tested set is given below each pair of bars; regulons represented by fewer than ten orthologues are hatched and are shown for completeness only. Enrichment relative to the background was assessed by one-sided Fisher’s exact test (*p < 0.05, **p < 0.01, ***p < 0.001, ****p < 0.0001; ns, not significant). **D** Relative expression of 6S RNA for *in vitro* grown *Y. pseudotuberculosis* at stress conditions. Data are presented as mean values ± SD from 3 sets (n=3) of *in vitro* samples, respectively. Pairwise comparisons to the control group (Ctrl) were performed using two sample t tests with Benjamini–Hochberg (BH) correction. Asterisks represent p-value thresholds: *** for p < 0.001, ** for p < 0.01, and * for p < 0.05; ns, non significant. Data for *in vitro* grownbacteria are from Avican et. al, 2021 (www.pathogenex.com). **E** Relative expression of 6S RNA under T3SS non-inducing (37°C) (left panel) and inducing (37°C, Ca^2+^ depletion) (right panel) condition in different oxygen concentrations for 3 hours measured with qPCR. Relative expression of 6S RNA is calculated according to Ct values normalized to that of *rpoB*. One-sample t tests were performed on log transformed relative expression values. Asterisks represent p-value thresholds: **** for p < 0.0001, *** for p < 0.001, ** for p < 0.01, and * for p < 0.05; ns, non significant. **F** Correlation between 6S RNA expression and total expression of pIBX virulence plasmid genes during infection. **G** The same correlation for bacteria grown in vitro at 26°C and at different time points of exposure to virulence-inducing conditions. Pearson correlation analysis was used in both **F**and **G**. **H** Relative 6S RNA expression under the *in vitro* conditions shown in **G**.

The heightened expression of 6S RNA in tissue bacteria, exceeding levels seen under any in vitro condition(23) indicates an *in vivo* specific expression pattern and suggests a key role for this ncRNA in tissue. This is supported by similarly elevated 6S RNA levels in *Y. pestis* recovered from infected mouse lungs compared to *in vitro* growth(30). In bacteria, 6S RNA binds the housekeeping RNA polymerase holoenzyme (RpoD; σ), inhibiting its binding to specific promoters and freeing RNA polymerase to associate with alternative sigma factors(31, 32). Given a potential role for 6S RNA in promoting survival in tissue, we hypothesized that bacteria entering this environment encounter multiple stressors requiring alternative sigma factors to induce genes for coping with stress. This assumption is supported by the increased expression of *rpoE* and *rpoH*, encoding alternative sigma factors involved in stress response, in tissue versus luminal bacteria (**Supplementary Table 2**). Sequestering RpoD via 6S RNA would thus facilitate transcriptional reprogramming upon host tissue entry. To verify this, we computationally predicted binding sites for RpoD, RpoS, and RpoE in *Y. pseudotuberculosis* as previously described for *E. coli*(33) incorporating experimentally validated transcriptional start sites and operon data(34) (**Supplementary Table S4**). Reanalysis of the 315 DEGs revealed that most genes upregulated in tissue lack a recognizable RpoD binding site (**Fig. 4B**), including 33 genes encoded on the virulence plasmid(35) 26 of which were significantly upregulated in tissue. To further validate the role of alternative sigma factors in tissue adaptation, we performed regulon enrichment analysis using experimentally supported sigma factor and gene interactions from RegulonDB, with orthologues assigned to *Y. pseudotuberculosis* by sequence alignment. Genes of the RpoH heat shock and RpoE envelope stress regulons were significantly overrepresented among genes upregulated in tissue, while the RpoS stationary phase regulon showed the same trend without reaching significance (9.6% against a 5.4% background, p = 0.065) (**Fig. 4C**). RpoE regulon enrichment is notable given elevated *rpoE* expression in tissue and the membrane perturbation expected to accompany T3SS assembly. RpoD regulated genes were also overrepresented; however, this regulon is dominated by housekeeping functions including translation and energy metabolism.

### 6S RNA expression tracks T3SS induction rather than oxygen

Consistent with its elevated expression in tissue, expression of 6S RNA was induced under the virulence-inducing condition and during microaerophilic growth at 26°C, among a set of ten different stresses applied separately (**Fig. 4D**). As oxygen level is critical for T3SS induction, we measured 6S RNA expression across a range of oxygen concentrations at 37°C, with and without virulence induction. Expression did not change significantly at any oxygen concentration (**Fig. 4E**), indicating that oxygen does not regulate 6S RNA expression. Instead, 6S RNA expression correlated with T3SS expression both *in vivo*, in tissue and lumen samples (**Fig. 4F**), and *in vitro*, across different durations of exposure to the T3SS-inducing condition (**Fig. 4G**). Moreover, 6S RNA was significantly increased after prolonged exposure to the virulence-inducing condition under aerobic conditions (**Fig. 4H**). Taken together, this analysis suggests a potential link between 6S RNA and T3SS induction in tissue-associated bacteria and under *in vitro* virulence inducing condition. To test whether 6S RNA is required for productive T3SS induction, we deleted *ssrS* and assessed Yop secretion. The mutant showed no growth defect (**Supplementary Fig. S4A**) and matched WT viability at all oxygen concentrations under both conditions (**Supplementary Fig. S5**), yet Yop secretion was reduced at 1% oxygen under inducing conditions (**Supplementary Fig. S6A and S7A, B**). This was not due to reduced synthesis: cellular Yop abundance was comparable between strains, apart from an increase in YopE at 5% oxygen (**Supplementary Fig. S8A, B**), and *yop* mRNA levels were unchanged except for *yopD* and *yopE*, which were higher in Δ*ssrS* at 1% oxygen under inducing conditions (**Supplementary Fig. S6B**). Loss of 6S RNA therefore affects a step downstream of effector synthesis, and the restriction to 1% oxygen indicates a conditional rather than a general requirement.

### 6S RNA restrains T3SS expression and limits surface immunogenicity *in vivo*

To define how 6S RNA shapes both the *Y. pseudotuberculosis in vivo* transcriptional programme and the host response to it, we orally infected four mice per strain with a low dose of WT or Δ*ssrS* and dissected the caecal luminal interface and tissue as before. We omitted antibody-based bacterial enrichment here, both because capture selects bacteria by surface antigen and could therefore differ between strains differing in envelope composition, and to retain host RNA for analysis from the same samples. Since WT bacterial transcriptomes were already available, only the Δ*ssrS* samples were sequenced, after depleting host RNA with dephoRNA(36) and bacterial rRNA with Jumpcode. Libraries were sequenced as described above, excluding one library with very few detected genes. Tissue and lumen replicates showed similar global expression density profiles (**Fig. 5A**), though lower overall coverage than WT infected samples reflected reduced mutant colonisation. PCA separated tissue and lumen samples, with tissue replicates clustering tightly (**Fig. 5B**) and showing strong global transcriptome correlation (**Fig. 5C**), whereas lower correlation among lumen replicates likely reflected zero inflation in gene detection (**Supplementary Fig. S9**). Differential expression analysis identified 279 significantly altered genes in Δ*ssrS* versus WT tissue bacteria (padj < 0.05; 190 upregulated, 89 downregulated; **Fig. 5D; Supplementary Table 5**), including 26 upregulated virulence plasmid genes associated with the T3SS. Consistently, qPCR confirmed increased *yopE* expression and detectable *yopH* transcripts in Δ*ssrS* tissue bacteria (which were undetectable in WT), while *yopD* levels remained unchanged (**Fig. 5E**). Virulence plasmid copy number was correspondingly elevated in Δ*ssrS* bacteria in tissue but not lumen (**Fig. 5F**), underlying the heightened T3SS expression. Because peak to trough ratios were indistinguishable between strains in both compartments (**Fig. 5G**), this reflects a specific increase in plasmid copy number rather than altered bacterial replication. Thus, 6S RNA restrains T3SS expression *in vivo*: its loss causes dysregulated T3SS overexpression rather than suppression, supporting our model that 6S RNA tunes T3SS induction to the level and timing required for productive tissue colonisation.

**Fig. 5.**
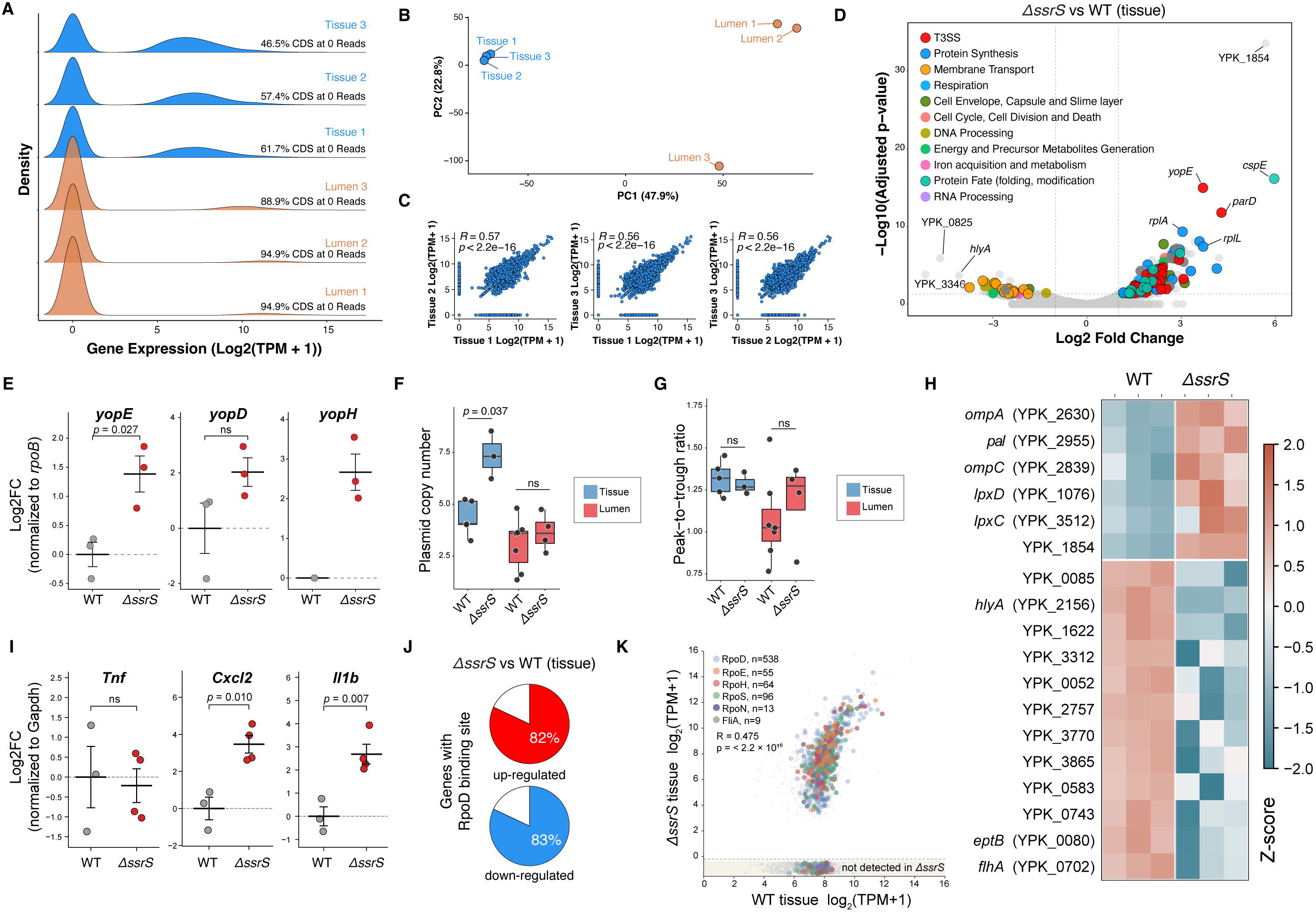
Loss of 6S RNA dysregulates the *in vivo* virulence programme, increases surface immunogenicity and shifts sigma factor regulon output. **A** Density distributions of gene expression (log (TPM + 1)) for Δ*ssrS* bacteria recovered from tissue (blue) and lumen (orange); the percentage of CDSs with zero mapped reads is indicated for each replicate. B Principal component analysis of global Δ*ssrS* transcriptomes. C Pairwise correlation of log (TPM + 1) values between tissue replicates; Spearman R and *p* values are indicated. D Volcano plot of differential expression in Δ*ssrS* versus WT tissue bacteria (log fold change against −log adjusted *p* value). Significantly altered genes (padj < 0.05) are coloured by PATRIC subsystem class; selected genes are labelled and dashed lines indicate significance and fold-change thresholds. **E** qPCR of *yopE*, *yopD* and *yopH* in WT and Δ*ssrS* bacteria recovered from caecal tissue, expressed as log fold change normalised to *rpoB*. **F** Virulence plasmid copy number in WT and Δ*ssrS* bacteria from caecal tissue and lumen, determined by droplet digital PCR. **G** Peak-to-trough ratio of genome coverage, reporting in vivo replication rate, for the same samples. **H** Z-score heatmap of surface-associated genes differentially expressed between WT and Δ*ssrS* tissue bacteria, calculated from library-size-normalised counts across three replicates per strain. The upper block shows genes up-regulated and the lower block genes down-regulated in Δ*ssrS*. **I** qPCR of host *Tnf*, *Cxcl2* and *Il1b* in caecal tissue from WT- and Δ*ssrS*-infected mice, expressed as log fold change normalised to *Gapdh*. **J** Proportion of genes up-regulated (red) and down-regulated (blue) in Δ*ssrS* tissue bacteria that carry a predicted RpoD binding site. **K** Mean TPM of each gene in WT tissue plotted against Δ*ssrS* tissue, log scale, for genes with combined mean TPM above 3. Genes assigned to a sigma factor regulon are coloured by regulon and genes without an assignment are grey. Genes not detected in Δ*ssrS* tissue (TPM = 0) are shown with vertical jitter in the shaded strip below the axis. The correlation shown is the mean of the nine individual Spearman correlations obtained by comparing each WT replicate against each Δ*ssrS* replicate, computed on log (TPM + 1) across all regulon genes. In **E** and **I**, points represent individual mice and bars show mean ± s.e.m.; *p* values are from Welch’s t-test applied to ΔCt values, with fold changes shown for display only. In **F** and **G**, boxes show median and interquartile range with points representing individual mice; *p* values are from Welch’s t-test comparing WT and Δ*ssrS* within each compartment. ns, not significant.

The most strongly induced gene in Δ*ssrS* was YPK_1854, encoding an LPP repeat protein with a lipoprotein leucine zipper domain (Pfam PF04728) typical of peptidoglycan anchored outer membrane lipoproteins, prompting us to examine surface immunogenicity. Surface associated genes upregulated in Δ*ssrS* included *ompA* (YPK_2630), *ompC* (YPK_2839), and *pal* (YPK_2955, peptidoglycan associated lipoprotein), which encode PAMPs that activate TLR2 and NF-κB dependent proinflammatory cytokine production(37–40). Upregulation of *lpxC* (YPK_3512) and *lpxD* (YPK_1076), encoding the first two rate limiting lipid A biosynthesis enzymes(41), further predicts increased surface LPS density and TLR4 stimulation. Conversely, genes downregulated in Δ*ssrS* mediate immune dampening. Four (YPK_0085, YPK_1622, YPK_2156, YPK_3312) encode filamentous haemagglutinin (FHA) family outer membrane proteins (**Fig. 5H**); in *Bordetella pertussis*, FHA engages complement receptor 3 on antigen presenting cells to induce IL-10 and suppress IL-12(42). A fifth, *eptB* (YPK_0080), encodes a phosphoethanolamine transferase whose orthologues modify lipid A to reduce TLR4 recognition(43, 44). To test this potential immune dampening directly, we measured expression of the proinflammatory mediators *Cxcl2*, *Il1b*, and *Tnf* by qPCR in caecal tissue from WT and Δ*ssrS* infected mice. Cxcl2 encodes the principal murine neutrophil chemoattractant(45) induced downstream of TLR2 and TLR4 via NF-κB(46), providing a direct readout of increased surface PAMP display. *Il1b* reports macrophage and inflammasome activation and potently induces *Cxcl2*(*47*), allowing us to test whether both arms of this axis respond. *Tnf* is a classical NF-κB target, but is also actively suppressed by the T3SS effector YopJ(48, 49), and is therefore informative for both host response and effector delivery. *Cxcl2* and *Il1b* were significantly elevated in Δ*ssrS*-infected tissue, whereas *Tnf* was unchanged (**Fig. 5i**). This selective induction is consistent with localised neutrophil recruitment driven by increased TLR2 ligand display, alongside continued YopJ-mediated suppression of TNF-α in bacteria overexpressing the T3SS. Together, these data indicate that 6S RNA coordinates two complementary layers of immune evasion during intestinal colonisation: suppression of TLR-activating surface PAMPs and maintenance of immunomodulatory surface proteins. Loss of 6S RNA collapses both layers simultaneously, predicting an elevated host inflammatory response in Δ*ssrS*-infected tissue.

### Loss of 6S RNA abolishes the RpoD-based partitioning of the tissue response

6S RNA sequesters RpoD-containing RNA polymerase, so its loss should release RpoD and erase the distinction between RpoD- and alternative sigma factor-dependent genes observed in WT tissue bacteria. Both predictions were met. Whereas genes up-regulated in WT tissue bacteria were largely devoid of recognisable RpoD binding sites (**Fig. 4B**), genes up- and down-regulated in Δ*ssrS* carried predicted RpoD sites at similar and high frequency (82% and 83%, respectively; **Fig. 5J**), indicating that the RpoD-based partitioning of the tissue response is lost in the mutant. Across all sigma factor regulons, WT and Δ*ssrS* tissue bacteria were less well correlated than replicates of the same genotype (R = 0.475, p < 2.2 × 10 ¹; **Fig. 5K**). Critically, Δ*ssrS* replicates agreed with one another as closely as WT replicates did (R = 0.66 to 0.69 versus 0.65 to 0.76 across the four regulons with sufficient orthologues; **Supplementary Fig. S10**), so the reduced correlation between genotypes reflects genuine divergence rather than noisier mutant libraries. Loss of 6S RNA thus releases RpoD-dependent transcription *in vivo* and abolishes the sigma factor partitioning that characterises the WT tissue response.

### Spatial localization of bacterial cells in tissue foci is important for adopting virulent and stress responsive states

Bacterial populations in host tissue are typically heterogeneous, allowing energetically costly virulence programmes to be partitioned among subpopulations(50–54). To characterise the cellular and spatial organisation of *Y. pseudotuberculosis* foci, infected caecal tissue was sectioned and stained with haematoxylin to locate the lymphoid aggregates containing bacterial foci (**Fig. 6A**). Combining immunohistochemistry against Ly6G with an RNAscope probe(55) for *Yersinia* 23S rRNA showed that the foci were surrounded by dense neutrophil infiltration (**Fig. 6B**), indicating a strong local immune response.To resolve bacterial subpopulations within these foci, we applied RNAscope probes for two virulence plasmid transcripts (*parE* and *lcrH*), 6S RNA, and fis as a marker of replicating cells (**Supplementary Table 6**). T3SS and 6S RNA signals were concentrated in bacteria near the tissue boundary, where neutrophils accumulated (**Fig. 6C**). Quantifying target signal only where it overlapped the 23S rRNA signal, across 43 tissue sections from two mice, revealed a heterogeneous population: 55% and 85% of cells expressed *parE* and *lcrH* respectively, around 50% expressed 6S RNA, and only ∼30% expressed *fis* (**Fig. 6D**). Approximately 70% of target-positive signal colocalised with 23S rRNA, confirming bacterial origin; the remainder likely reflects low ribosomal content in quiescent cells or limited probe access within tightly packed foci (**Supplementary Fig. S11A, B**). Mapping signal intensity against distance from the foci–tissue boundary showed that 23S rRNA dominated in the focus interior, whereas *parE* and 6S RNA were comparatively stronger at the boundary, giving a red-to-green ratio that peaked closest to host tissue (**Fig. 6E, F**). *lcrH* and *parE* were rarely detected on the luminal-facing side, whereas *fis* showed no spatial bias. *Y. pseudotuberculosis* therefore adopts spatially distinct transcriptional states within tissue foci, with virulent and stress-adapted cells at the boundary facing host immune cells and replicating cells distributed throughout.

**Fig. 6.**
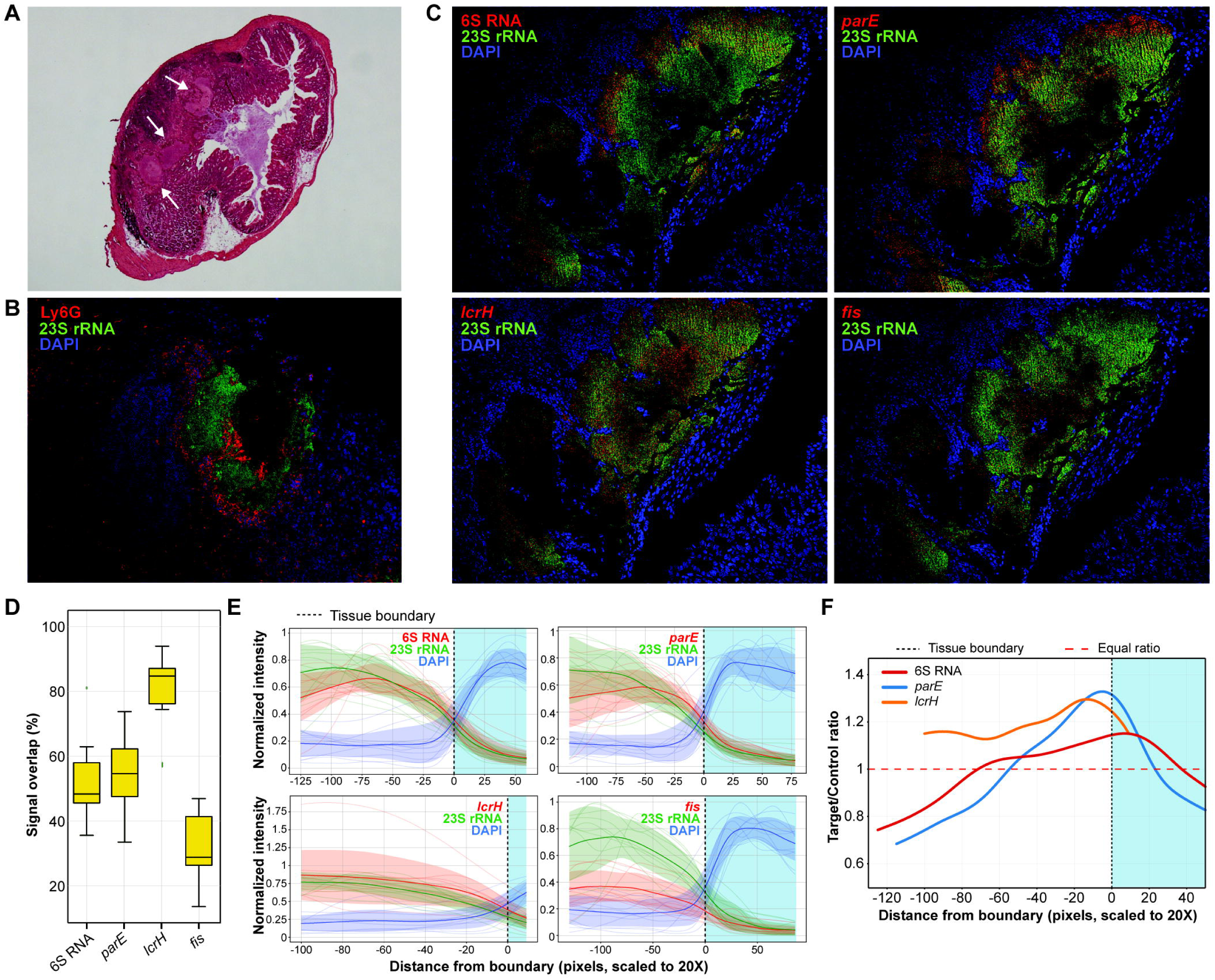
Bacterial foci in cecal lymphoid tissue are surrounded by neutrophils and display spatially organized *Yersinia* subpopulations with heterogeneous expression of 6S RNA and T3SS genes. **A** Hematoxylin staining of mouse cecum infected for 2.5 days with WT *Y. pseudotuberculosis*. Bacterial foci in lymphoid tissue are indicated with white arrows. B Combined immunohistochemistry with RNAscope Assay on infected cecal tissue section. Sections were stained with anti-Ly6G targeting neutrophils (red signal), DAPI (blue signal), and an RNAscope probe targeting *Y. pseudotuberculosis* 23S rRNA (green signal). C Representative tissue sections of a bacterial foci in cecal tissue stained with DAPI (blue signal), and RNAScope Assay probes targeting 6S RNA (red signal, top left panel), *parE* (red signal, top right panel), *lcrH* (red signal, bottom left panel), *fis* (bottom, right panel), and 23S rRNA (green signal, all panels). D Box plot showing percentages of 23S rRNA signals overlapping with 6S RNA, *parE*, *lcrH,* and *fis* signals in multiple bacterial foci of separate tissue section stainings (6S RNA; n=11, *fis;* n=11, *lcrH;* n=10, *parE;* n=11) from two individual mice (n=2) E Line plot of normalized red (target gene), green (23S rRNA), and blue (nuclei) signal intensities and their distance to the bacterial foci-tissue boundary. The vertical black dashed line indicates the foci-tissue boundary and light-blue background in the plots indicate tissue region. Normalization was performed as follows: the blue channel was individually normalized for each sample using min–max scaling, i.e. (signal − min) / (max − min + ε), where ε is a small constant added to avoid division by zero. The red and green channels were normalized using the min and max values of the green channel. This approach allows the usage of the green channel as a control reference, making it easier to detect relative variation between RNA targets in the red channel. The initial line plots were smoothed by applying a Gaussian filter to the raw intensity profiles. F Average of target (red signal from 6S RNA, *parE*, or *lcrH*) and control (green signal from 23S rRNA) ratio in relation to the distance to foci-tissue boundary of all analysed foci in different tissue sections.

### 6S RNA accelerates tissue colonization

To test the importance of 6S RNA for pathogenicity *in vivo*, we performed competitive infection with a 1:1 mixture of tdTomato-expressing WT and GFP-expressing Δ*ssrS* and counted CFUs from caecal tissue and lumen at 2.5 days. A control co-infection of the two differently labelled WT strains gave comparable loads, confirming that the reporters do not affect infection (**Supplementary Fig. S12**). Although bacterial loads varied between animals, Δ*ssrS* colonised the lumen as efficiently as WT while its recovery from tissue was significantly reduced, with the competitive index falling from 0.79 in the lumen to 0.32 in tissue (**Fig. 7A**). 6S RNA therefore contributes to efficient colonisation of the tissue compartment. In single-strain infections with unlabelled WT and Δ*ssrS*, CFUs were quantified in whole caecal content and in gentamicin-treated lymphoid tissue biopsies, which restricts counts to tissue-associated bacteria. At 1 dpi luminal loads exceeded tissue loads for both strains, indicating that translocation had not yet occurred (**Fig. 7B**). By 2 dpi, the timepoint at which the RNA-seq samples were collected, the two compartments were equivalent for WT whereas luminal loads remained significantly higher for Δ*ssrS*, indicating delayed translocation. Since Δ*ssrS* infection elevated *Cxcl2* and *Il1b* in tissue (**Fig. 5I**), we asked whether this heightened response extended to the mucosal surface and measured luminal lipocalin-2, a marker of intestinal inflammation released by neutrophils and epithelium. Lipocalin-2 trended higher in Δ*ssrS-*than WT-infected mice at 2 dpi and was significantly elevated at 3 dpi (**Fig. 7C**). It also correlated with bacterial load in Δ*ssrS*-infected mice (r = 0.59, *p* = 0.017) but not in WT (r = 0.22, *p* = 0.42; **Fig. 7D**), indicating that WT bacteria uncouple their burden from the intensity of the host response and that this uncoupling is lost without 6S RNA. Delayed tissue colonisation by Δ*ssrS* is therefore accompanied by a stronger innate response per bacterium, consistent with increased immune visibility.

**Fig. 7.**
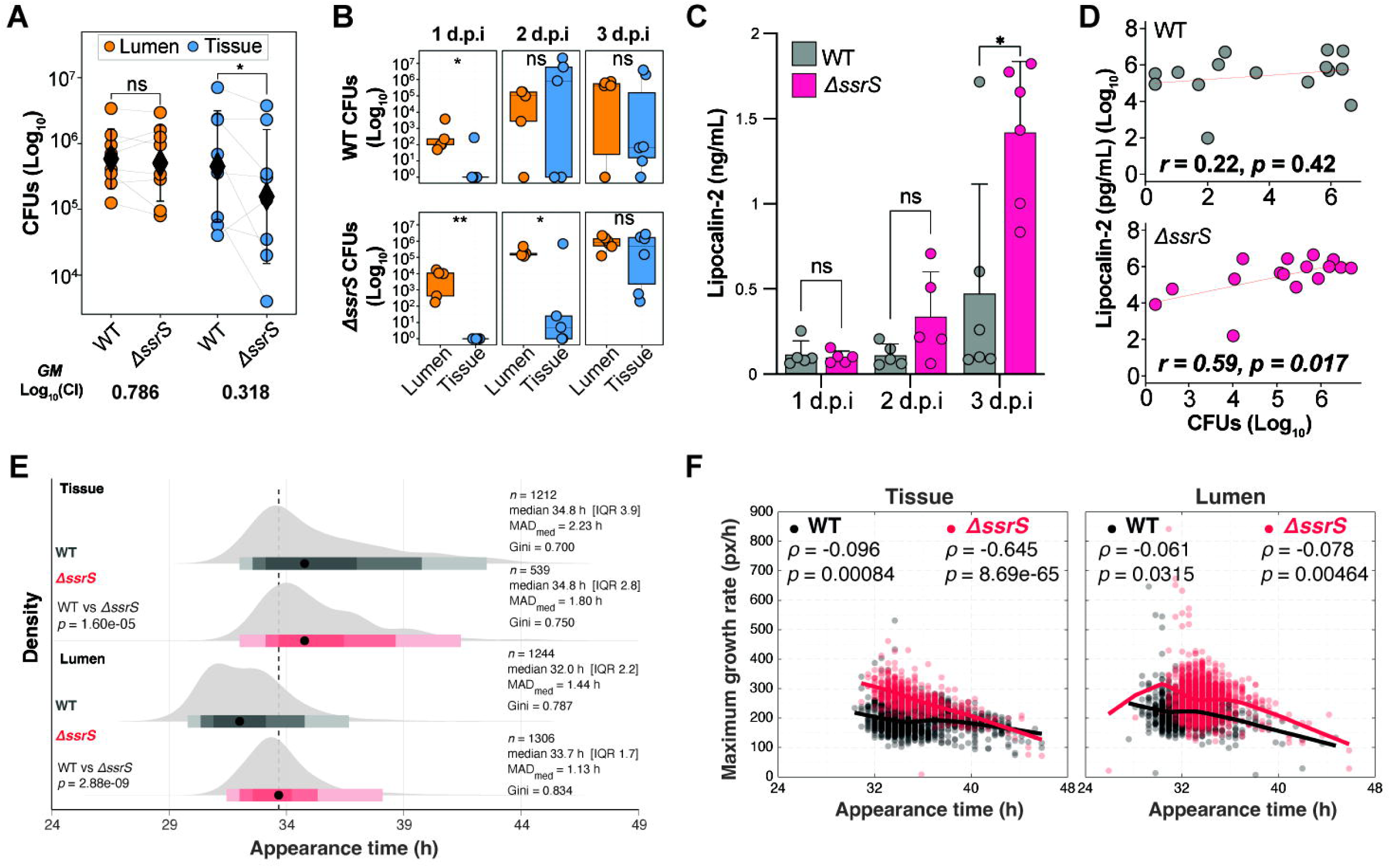
6S RNA promotes timely tissue colonisation, limits host inflammation and shapes the physiological diversity of tissue-associated bacteria. **A** CFUs recovered from cecal tissue and lumen compartments of mice (n=8) co-infected with 1:1 mixture of *Y. pseudotuberculosis* WT:*tdTomato* and Δ*ssrS*:*gfp* strains at 2.5 dpi. The two strains were distinguished with fluorescent signal by tdTomato expression in WT strain (WT:*tdTomato*) and by GFP expression in Δ*ssrS* strain (Δ*ssrS*:*gfp*). The competitive index was calculated as the ratio of Δ*ssrS* CFUs to WT CFUs, and the geometric mean CI was reported. The geometric mean is indicated by a diamond symbol, and the standard error of the mean (SEM) is shown as error bars. Statistical significance was calculated with Wilcoxon signed rank test. B CFUs recovered from single strain infection of mice with *Y. pseudotuberculosis* WT (n=5) and Δ*ssrS* (n=5) strains. CFUs recovered from cecal lumen and tissue at 1, 2 and 3 dpi. Data are presented as Log-transformed CFU counts for each compartment. Comparisons between paired lumen and tissue samples were performed using a paired t test applied to Log-transformed CFU values. C Lipocalin−2 levels measured with ELISA from cecal lumen of mice in B. Multiple comparisons and statistical significance of Lipocalin-2 concentrations (C) were calculated by two-factor ANOVA test (main effects and repeated measures were considered) in Prism 10. D Correlation analysis between Lipocalin 2 concentrations and bacterial load. Pearson correlation analysis was applied to log_10_ transformed Lipocalin 2 concentrations and log_10_ transformed CFU counts. E-F Colonies recovered from the tissue and lumen of WT and Δ*ssrS* were plated on Luria-Bertani agar, incubated at room temperature for 20h, and imaged in the HeteroTyper system for 28h at 30 minutes intervals. WT is shown in dark gray and Δ*ssrS* in red. E Colony appearance time shown as density ridgeline plots. Black dots below each filled area represent the group median, and bars show the central 50% (interquartile range, IQR; darkest), 80%, and 95% (lightest) of colonies. The vertical dashed line marks the median of all groups. Significance was tested using the Brown-Forsythe test followed by ANOVA and corrected using Benjamini-Hochberg (BH). The number of colonies (n) and mean absolute deviation from the group median (MAD_med_) are reported per group. F Colony appearance time versus maximum growth rate (px/h), estimated by fitting a Gompertz model to colony area over time, for colonies from tissue (left) and lumen (right). Each dot represents an individual colony. Points and locally estimated scatterplot smoothing (LOESS) curves are colored by strain. Spearman correlation coefficients (ρ) and *p*-values (*p*) are shown for each strain and infection site. Appearance time is represented in hours (h), while the colony area was measured in pixels (px).

### 6S RNA shapes phenotypic heterogeneity of tissue-associated bacteria

To test whether the transcriptional and inflammatory differences described above are reflected in the physiological states of individual bacteria, we analysed colonies recovered from infected tissue and lumen using HeteroTyper, a time-lapse colony imaging platform(56). Colony appearance time served as a proxy for lag, and its dispersion within a population reports the diversity of physiological states present at recovery(57–59). Appearance times were more dispersed in tissue-than lumen-derived colonies for both strains (**Fig. 7E**), consistent with tissue being the more stressful environment. WT tissue colonies were more dispersed than Δ*ssrS* colonies, quantified as the median absolute deviation from the median (MAD_med), despite identical median appearance times of 34.8 h, indicating that 6S RNA contributes to the generation of heterogeneous physiological states. Appearance time also predicted subsequent growth, but only in the mutant. In WT the two were effectively independent in both compartments (tissue ρ = −0.096; **Fig. 7F**), whereas Δ*ssrS* colonies from tissue were strongly anticorrelated (ρ = −0.645); Δ*ssrS* lumen colonies behaved as WT did (ρ = −0.078). This follows from the role of 6S RNA in recovery from growth arrest: it sequesters RpoD-containing RNA polymerase during stationary phase in a form that pRNA synthesis releases upon nutrient upshift, so WT cells restart from a pre-formed reserve however long they remained quiescent. Without it, RpoD-dependent transcription must be rebuilt and the duration of quiescence sets both lag and growth rate, which matters only where bacteria span a broad range of quiescent states (**Fig. 7E**). The phenotype was reproduced *in vitro* without any host, the Δ*ssrS* correlation reaching ρ = −0.593 against ρ = −0.301 for WT after 72 h in stationary phase (**Supplementary Fig. S13B**). The mutant also appeared later than WT at every growth phase and under acidic, iron-limiting and osmotic stress, though not oxidative stress (**Supplementary Fig. S13A, S14A**). The inverse relationship was present in both strains under every condition but stronger in the mutant only under iron limitation and osmotic stress (**Supplementary Fig. S14B**), the two in which WT coupling was weakest, as in tissue. 6S RNA therefore supports efficient recovery from growth arrest across a range of stresses, though not universally.

## Discussion

This study provides the first evidence of a pronounced, spatially restricted induction of 6S RNA in a bacterial pathogen during host infection. Previous transcriptomic surveys, including the PATHOgenex RNA atlas(23), catalogued non-coding RNA expression across pathogens and stress conditions without reporting *in vivo* 6S RNA induction of this magnitude, and the host-niche specificity described for *Rickettsia conorii*(60) was confined to cultured cells. Infection phenotypes have been reported for 6S RNA in *Legionella pneumophila*(61) and *Salmonella* Typhimurium(62), but in neither case was the RNA connected to a virulence programme or to increased host recognition. Our data place 6S RNA at the transition from lumen to tissue, where it restrains type III secretion and envelope immunogenicity.

An important consequence of the spatial restriction we observe is that bulk transcriptomic measurements necessarily understate induction within responding cells. If only half the population induces 6S RNA, the fold change recovered from bulk RNA-seq is diluted by the non-expressing fraction, and the induction experienced by individual responding bacteria is correspondingly greater. Partitioning of costly virulence programmes across a population is well documented for the T3SS and is thought to distribute the metabolic burden of virulence while maintaining a replicating subpopulation(50, 63), and heterogeneous pathogen gene expression drives heterogeneous host responses in turn(64). Our data indicate that 6S RNA induction is partitioned in the same way, and that the regulatory shift we describe is concentrated in the subpopulation directly engaging host immune cells.

That 6S RNA acts as a restraint rather than a switch is reinforced by a second candidate restraint in the same dataset. The nucleoid-associated protein H-NS (YPK_2074) and the integration host factor subunit IHF-α (YPK_1826) were both induced in tissue-located bacteria. H-NS silences horizontally acquired, AT-rich sequences including the *Yersinia* virulence plasmid, and the H-NS/YmoA complex represses the T3SS in a manner that permits IscR-dependent regulation(17). Elevated H-NS during T3SS induction is therefore unlikely to reflect simple de-repression, and more plausibly provides a counterweight holding T3SS expression within a productive range. RpoE may constitute a third, responding to membrane stress and induced upon virulence induction *in vitro*(23); whether its induction here reflects envelope perturbation by the environment or by the secretion apparatus itself remains unresolved. Together these restraints may explain why loss of 6S RNA produces overexpression rather than loss of the T3SS, and why that overexpression is nonetheless uncoupled from effective secretion.

The enrichment of both 6S RNA and T3SS transcripts at the neutrophil interface in wild-type foci, alongside T3SS overexpression in the Δ*ssrS* mutant, points to negative feedback on T3SS output. In this model, host-interface cues induce the T3SS, while sustained exposure to those cues also induces 6S RNA, which then caps T3SS output. Consistent with a delayed brake rather than a parallel input, 6S RNA accumulated only after prolonged virulence induction in vitro (**Fig. 4H**) and was unaffected by oxygen (**Fig. 4E**), one of the cues driving T3SS expression. Removing this restraint increased virulence plasmid copy number and T3SS transcription. Because each RNAscope target was imaged in a separate section, these data establish co-enrichment of 6S RNA and T3SS transcripts within the same region of the focus rather than co-expression within individual cells, and the ordering of induction is inferred from in vitro kinetics rather than measured in tissue. Multiplexed detection of 6S RNA and T3SS transcripts within individual bacteria, combined with a time course of tissue infection, would establish whether 6S RNA indeed rises after T3SS in the cells engaging host immune cells.

The host response tracks this failure of restraint. *Yersinia* uses the T3SS to suppress innate signalling(12, 65, 66), and in WT infection luminal lipocalin-2 was unrelated to bacterial burden, whereas in the mutant it scaled with burden. Wild-type bacteria therefore appear to uncouple how many organisms are present from how strongly the host responds, and this uncoupling is lost without 6S RNA.

Single-colony analysis provided an independent view of the same defect. Appearance time reports the physiological state of individual cells at recovery, since prior stress delays resumption of growth(58, 59). Appearance time and subsequent growth rate were uncoupled in wild-type but strongly anticorrelated in Δ*ssrS* bacteria from tissue, and this was reproduced in culture, most strongly after 72 h in stationary phase. 6S RNA sequesters RpoD-containing RNA polymerase during growth arrest and releases it upon nutrient upshift(31, 32, 67), so wild-type cells resume RpoD-dependent transcription from a preformed reserve irrespective of how long they remained quiescent, whereas in the mutant no such reserve exists and cells emerging from deeper quiescence restart later and grow more slowly thereafter. The mutant was correspondingly delayed under acid, iron-limiting and osmotic stress, though not under oxidative stress, indicating that this requirement is broad rather than specific to the tissue environment.

Tissue-located bacteria encounter several cues simultaneously, and our transcriptome indicates at least iron limitation, oxidative stress and nutrient restriction acting together, whereas each *in vitro* condition isolates one. The coupling between appearance time and growth rate was stronger in the mutant under iron limitation and osmotic stress but not under acid or oxidative stress, and these two conditions are among those most plausibly combined in tissue. Single stressors may therefore under-represent the composite state the bacterium actually experiences, and the apparent condition dependence may reflect the limits of the *in vitro* panel rather than a genuinely restricted requirement for 6S RNA. The same consideration applies to oxygen: although oxygen tension and iron availability regulate the Ysc T3SS(13, 68), *lcrF*, its downstream reporter *rpsJ*(69) and *iscR*(70) were expressed comparably in both compartments, indicating that oxygen alone does not account for the enhanced T3SS and 6S RNA activity on tissue entry.

The regulatory programme described here would have been invisible in culture. Transcriptomes of tissue-located bacteria separated cleanly from those of luminal bacteria only millimetres away, and neither resembled any of the in vitro conditions tested (**Supplementary Fig. S1**), with 6S RNA reaching levels that no single stress reproduced. Recovering such programmes directly from infected tissue remains technically demanding, because pathogen transcripts constitute a vanishing fraction of the RNA in an infected biopsy, and this bottleneck is still being actively addressed: capture-based enrichment has recently been developed to retrieve *Mycobacterium tuberculosis* mRNA from host samples(71), and *in vivo* dual RNA-seq of extracellular pathogens is only now reaching the sensitivity needed to resolve bacterial effector programmes during barrier crossing(72). Antibody-based enrichment of *Yersinia* cells prior to RNA extraction raised bacterial mRNA from 0.002% to 0.082% of reads, a 41-fold gain yielding near-complete coding-sequence coverage. Full coverage, rather than depth alone, is what made this study possible: our earlier work on the same infection model could not reach complete transcriptomes(13), and identifying 6S RNA as the most strongly induced transcript depends on having measured the rest of the transcriptome alongside it.

A regulator that appears dispensable in the laboratory becomes, in tissue, the element that holds virulence output and immune visibility in balance, and it does so in the small subpopulation that meets the host directly.

## METHODS

Detailed procedures are provided in SI Appendix, Supplementary Materials and Methods.

### Bacterial strains and growth

The bioluminescent, kanamycin-resistant *Y. pseudotuberculosis* strain YPIII/pIBX was used as WT strain in this study. The strain YPIII has long served as a classic model system for studying *Y. pseudotuberculosis* pathogenesis and has played a key role in the discovery and analysis of Yops(73–78) and used for persistent mouse model infections(11, 13). For *in vitro* growth, *Y. pseudotuberculosis* WT and mutant strains were grown in LB with 50 μg/ml kanamycin supplementation at 26°C. For T3SS induction conditions, overnight cultures were subcultured with 1/100 dilutions, grown to OD_600_:0.3 at 26°C and thereafter shifted to 37°C with addition of 20 mM MgCl_2_ and 10 mM EGTA for three hours. Same protocol, except addition of 20 mM MgCl_2_ and 10 mM EGTA, was employed for T3SS non-inducing condition at 37°C. For T3SS induction and non-inducing conditions under hypoxia and different oxygen levels (0%, 0.1%, 1%, 5% and 10%), bacterial cultures were grown overnight and subcultured with 1/100 dilutions, grown to OD_600_:0.3 at 26°C under aerobic conditions and cells were pelleted with centrifugation at 8000 *g* for 2 minutes. After removal of supernatants, pellets were moved to anaerobic chamber (Whitley A35 Workstation, Don Whitley Scientific Limited, England) set to desired oxygen level one day prior to induction. Pellets were resuspended in 24 hours pre-incubated LB in the chamber and supplemented with 20 mM MgCl_2_ and 10 mM EGTA for T3SS induction condition. Desired gas configuration was done according to manufacturer’s manual and recommendations.

### Ethical statement and approval for mouse studies

Mice were housed in the infection unit at the Umeå Centre for Comparative Biology (UCCB) in accordance with the Swedish National Board for Laboratory Animals guidelines. All animal procedures were approved by the Animal Ethics Committee of Umeå University (Dnr A54/2015). During arrival, prior to experiments, the mice were acclimated for one week.

### Mouse infection and *in vivo* imaging

Methods for low dose infection are similar to Avican et al. (2015) with few modifications. In brief, eight-week-old female FVB/N mice (Charles River Laboratories, Inc.) were deprived of food and water for 16 hours and subsequently provided 0.5×10^6^ CFU/ml *Y. pseudotuberculosis* YPIII or Δ*ssrS* mutant strain resuspended in tap water supplemented with 150 mM NaCl, and administered via their drinking water for 6 hours. Final infection dose was determined via viable counting on LB agar plates with kanamycin (50 µg/ml) and the drinking volume consumed after 6 hours, divided by the number of mice in each cage. For co-infection experiment, WT:*tdTomato* and Δ*ssrS:gfp* strains were mixed with 1:1 ratio and resuspended in tap water supplemented with 150 mM NaCl and administered to mice via their drinking water for 6 hours. Final infection dose for each strain was determined via viable counting of colonies via fluorescent stereo microscope on LB agar plates with kanamycin (50 µg/ml) and ampicillin (100 µg/ml) the drinking volume consumed after 6 hours, divided by the number of mice in each cage. In accordance with the guidelines for Laboratory Animals, mice were inspected frequently for signs of infection and clinical symptoms that may show low possibility of recovery, minimizing suffering in the sick animals. Infections were also monitored using an IVIS spectrum (Caliper LifeSciences, Inc.). IVIS analysis of the mice was performed under anaesthesia using an XGI-8 gas anesthesia system (Caliper LifeSciences, Inc). Oxygen mixed with 2.5% IsoFloVet (Orion Pharma, Abbott Laboratories Ltd, Great Britain) was used to induce anaesthesia and maintained using oxygen with 0.5% isoflurane during the duration of the imaging. Images were then acquired and analyzed using Living Image software (Caliper LifeSciences, Inc.).

### Bacterial cell enrichment and total RNA isolation from cecal tissue and luminal interface

To enable spatial tissue separation, mice with clear bioluminescent signals were subjected to careful surgical separation of the caecal aggregate and are subsequently identified as our deeper tissue. The luminal side of the aggregate, which was exposed after dissection of tissue samples, was also sampled and placed in separate Dispomix tubes (Medic Tools AG, Switzerland) with 1mL cell RNAprotect (Qiagen, USA) to stabilize the RNA. Tissue homogenization in the tubes with a Dispomix Drive using homogenization program 9 (Medic Tools AG, Switzerland). Samples were quickly spinned down at low speed (350 *g*) to allow the large particulates to settle. The supernatant was transferred to new tubes and added with 20 µl of antibody-coupled beads prepared previously as above. Sample and beads are transported together and performed according to the Dynabeads anti-*Salmonella*/anti-*E. coli* (Invitrogen, Lifescience Technologies, CA, USA) protocols. The mix of sample and beads was incubated for 10 minutes at room temperature with gentle continuous inversion to prevent the beads from settling. Tubes were then inserted in a magnetic rack (DynaMag^TM^-2) and allowed to stand for three minutes (or until clear) and carefully tilted and inverted to allow all of the anti-*Y. pseudotuberculosis* coupled Dynabeads to be captured on the side of the magnet for maximum recovery. The supernatant was carefully removed, and magnetized Dynabeads washed with 1mL buffer twice. After the final wash, the anti-*Y. pseudotuberculosis* coupled Dynabeads with enriched *Y. pseudotuberculosis* cells were directly resuspended in 500 µl TRIzol^TM^ reagent and kept in −80°C until extraction. Total RNA extraction was done using TRIzol^TM^, using a TRIzol^TM^ column and using the TRIzol^TM^ DNA extraction protocol according to manufacturer’s instructions (Invitrogen, Lifescience Technologies, CA, USA). Quality of extracted RNA was then checked using agarose gel electrophoresis to make sure that there is no gDNA contamination and degradation of the extracted total RNA.

### RNA-seq library preparation and sequencing

Total RNA extracts were enriched in *Y. pseudotuberculosis* mRNAs using MICROB*Enrich* Kit (Ambion), which removes 18S and 26S rRNAs and polyadenylated (polyA) mRNAs according to the manufacturer’s instructions. RNA pools extracted from these samples were then further depleted of ribosomal RNAs using RiboZero Epidemiology Gold kit (Epicentre) and proceeded following preparation of strand-specific libraries using ScriptSeq V2 kit (Epicentre) according to ScriptSeq Complete Kit (Bacteria), with minor modifications in the 2-step cleanup to optimize RNA concentration output. Libraries were run on an Illumina Novaseq 6000 (SciLifelab, Sweden) for a sequencing depth of approximately 100M to 1B paired-end reads of 150 nt read length.

### RNA-seq and differential expression analysis

Adapter and quality trimming was done using Cutadapt v2.5 on 150-nt-long paired-end Illumina Novaseq 6000 reads. Mapping was done on *Y. pseudotuberculosis* YPIII (NC_010465), and the pYV plasmid (NC_006153) from NCBI were used as reference genomes for mapping using CLC Genomics Workbench. Annotations for rRNA and tRNA were removed from the reference genomes prior to mapping to minimize bias that may be introduced from the rRNA depletion procedures. Preliminary analysis of total mRNA read counts and normalized expression values for each gene were performed using RUVseq to determine the extent of potential batch and replicate effects from the *in vivo* samples(79–81). The sample types that can have the least unwanted variance were determined and subjected for differential expression (DE) analysis using edgeR(82, 83) to scale for total number of reads in Counts per Million (CPM), which was used as input for analysis using DEseq2 median of ratios method(84) to account for differences in sequencing depth and RNA composition that can skew the analysis.

Differential expression between wild-type (WT) and Δ*ssrS Y. pseudotuberculosis* in the cecal tissue compartment was assessed using DESeq2 (v1.40) implemented in R (v4.3). Raw count matrices were derived from bacteria recovered from the cecal tissue of infected mice. Prior to differential expression testing, low-confidence genes were removed using the following criteria: genes must have a non-zero count in all three WT libraries (ensuring the gene is robustly expressed under WT *in vivo* conditions), in at least two of the three retained Δ*ssrS* libraries (allowing for structural zeros attributable to low bacterial loads rather than true absence), and a total raw count of ≥ 10 across all six libraries. This filter was applied before size factor estimation to avoid fitting dispersion curves to near-empty rows. Size factors were estimated using the poscounts method (sfType = “poscounts”), which computes normalization constants from the geometric mean of non-zero counts per gene and is robust to excess structural zeros. Standard median-of-ratios size factors were inappropriate here because the Δ*ssrS* libraries exhibited pronounced zero-inflation (approximately 50% of filtered genes undetected in at least one mutant library, versus near-complete detection in WT), a condition that systematically biases the median-of-ratios estimator. Dispersion was estimated using the standard DESeq2 shrinkage pipeline. Cook’s distance-based outlier replacement was disabled (cooksCutoff = FALSE; minReplicatesForReplace = Inf) because outlier detection and replacement are poorly powered with n = 3 replicates and can inflate false positives at low replicate numbers. Differential expression was estimated by the Wald test using the contrast Δ*ssrS* versus WT; a positive log2 fold change therefore denotes higher expression in Δ*ssrS*. P values were adjusted for multiple comparisons using the Benjamini– Hochberg (BH) procedure; genes with an adjusted p value (padj) < 0.05 were considered statistically significant. To verify that differentially expressed genes were not an artifact of the depth imbalance between WT and Δ*ssrS* libraries, a depth-matched sensitivity analysis was performed. All six libraries were rarefied to 13,000 reads (the depth of the shallowest retained Δ*ssrS* library) by multinomial subsampling without replacement (rmultinom, set.seed(1)). DESeq2 analysis was repeated on the rarefied matrix using identical parameters. Concordance between the full-depth and depth-matched analyses was used to confirm that the directional shifts reported below reflect genuine transcriptional differences rather than sequencing-depth artefacts.

## Data availability

All sequencing data and processed data have been deposited in the GEO database with accession code GSE300615.

## Code availability

HeteroTyper pipeline used for image processing and downstream analyses is available as open-source software on GitHub (https://github.com/avicanlab/HeteroTyper) and Zenodo (https://doi.org/10.5281/zenodo.23951321). The comprehensive documentation is hosted on ReadTheDocs (https://heterotyper.readthedocs.io/). The infection and *in vitro* growth and stress data generated in this study are deposited on Zenodo (https://doi.org/10.5281/zenodo.23952201).

## Acknowledgement

This work was supported by Swedish Research Council (No. 2021-02466), Kempestiftelserna (JCK22-0017), and the Medical Faculty at Umeå University (FS 2.1.6-281-22) to K. Avican, by Swedish Research Council Excellence Center grant (No. 2022-06543) for the Center for Modeling Adaptive Mechanisms in Living Systems Under Stress to K. Avican and M. Fällman, by Swedish Research Council (No. 2022-00692) and Insamlingsstiftelsen, Medical Faculty at Umeå University to M. Fällman. We acknowledge µNordic Single Cell Hub (µNiSCH) and Small animal research imaging facility (SARIF) at Umeå University. We thank Niklas Habig for his assistance in qPCR experiments in this study. We thank Scilifelab and National Genomic Infrastructure (NGI) for sequencing RNA-seq libraries.

## Author contributions

OS, VDV, UA, SGS, TD, SB, AF, JG, NH, FM, KN, AN, HW, MF, and KA conceived, designed, and performed the experiments and analyzed the data. OS, MF, and KA wrote the manuscript with input from VDV, UA, SGS, TD, SB, AF, JG, NH, FM, KN, AN, and HW.

