## Supplemental Information for "6S RNA tunes virulence, surface immunogenicity, and phenotypic heterogeneity in tissue-invading *Yersinia*"

**Supplementary Information**

**Supplementary Materials and Methods**

**Gentamicin Treatment Assay**

Tissue biopsy samples were processed according to established gentamicin treatment protocol(1). Briefly, each isolated cecum was opened content removed, rinsed with PBS, and transferred into a sterile microcentrifuge tube and incubated in 1 mL of gentamicin solution (400 µg/mL in PBS) for 30 minutes at room temperature to eliminate extracellular bacteria. Following antibiotic treatment, samples were washed nine times with 2 mL sterile PBS to ensure complete removal of residual gentamicin. The treated tissues were then homogenized and the resulting homogenates were plated onto LB agar supplemented with kanamycin (50 µg/mL) for selective recovery of intracellular bacteria.

**Preparation of Dynabeads coupled with polyclonal *Yersinia* antibody**

The manufacturer protocol for Dynabead Antibody Coupling Kit (Invitrogen^TM^, Life Technologies, CA, USA) was used with modifications. In detail, dry beads were resuspended with the correct volume of purified anti-*Yptb* polyclonal antibodies(2) (approximately 2mg/µl) and buffer A.

### ***Y. pseudotuberculosis* sigma factors binding site determination**

500 nucleotide upstream from transcription start site of a gene or operon were considered for sigma factor binding sites prediction. DNA binding motifs of sigma factors were collected from RegulonDB(3). Motifs which were experimentally validated in *E. coli* were considered and  converted in to Position weight matrix (PWM). PWM for each sigma factor motif were searched on the upstream sequences by an in-house built computer program written in Perl language. Prediction score accuracy was based on finding a motif within the upstream sequence above genomic noise. Genomic noise was predicted from the equal length of DNA taken randomly from the genome. Majority of the TSS of genes and operons were collected from Nuss et al., 2015 and unreported TSS were predicted insilico by PromoTech tool (https://github.com/BioinformaticsLabAtMUN/PromoTech).

**Sigma factor regulon enrichment analysis**

Sigma factor–gene regulatory interactions for *Escherichia coli* K-12 were retrieved from RegulonDB v14.5(4) using the NetworkSigmaGene dataset, which contains experimentally supported interactions between sigma factors and their regulated genes together with confidence levels. Six sigma factors were analysed: σ⁷⁰ (RpoD, housekeeping), σ²⁴ (RpoE, extracytoplasmic stress), σ³² (RpoH, heat shock), σ³⁸ (RpoS, stationary phase), σ⁵⁴ (RpoN, nitrogen) and σ²⁸ (FliA, flagellar). σ¹⁹ (FecI) was excluded because RegulonDB lists only five target genes, none with a *Y. pseudotuberculosis* orthologue. Analyses were restricted to interactions annotated as Confirmed or Strong evidence. Because RegulonDB gene sets are defined for *E. coli* K-12, regulon members were mapped to *Y. pseudotuberculosis* YPIII by sequence-based orthology. The YPIII proteome was derived by translating all annotated CDS features (NCBI translation table 11) from the genome sequence and GFF3 annotation, yielding 4,269 protein sequences. Reference *E. coli* K-12 proteins were obtained from UniProt proteome UP000000625 (reviewed entries; 3,169 sequences with assigned gene names), of which 1,629 corresponded to RegulonDB sigma factor targets. Pairwise local alignments were computed using the Smith–Waterman algorithm (BLOSUM62, gap opening penalty 11, gap extension penalty 1) with the SIMD-accelerated Opal implementation via pyOpal v0.7.3. Each *E. coli* regulon protein was aligned against the complete YPIII proteome and the highest-scoring hit retained; each such YPIII protein was then aligned against the complete *E. coli* proteome. Pairs that were mutual best hits in both directions were retained as orthologues, yielding 1,175 reciprocal pairs (72% of queries; median amino acid identity 79%, median query coverage 100%). Orthologues were further filtered at ≥60% amino acid identity and ≥60% query coverage, giving final regulon sizes of 538 (σ⁷⁰), 96 (σ³⁸), 64 (σ³²), 55 (σ²⁴), 13 (σ⁵⁴) and 9 (σ²⁸) *Y. pseudotuberculosis* genes. Enrichment was assessed separately within each differential expression comparison using one-sided Fisher’s exact tests.

**Construction of *ssrS* deletion mutant**

To create *ssrS* deletion mutant in *Y. pseudotuberculosis*, DNA fragments flanking *ssrS* gene, aimed to be deleted, were amplified. Using In-fusionHD cloning kit (Clontech) the fragment was cloned into the *Sac*I/*Xho*I-digested suicide vector pDM4 which were then transformed into *E. coli* *S17-1λpir*. This strain was used as donor strain in conjugational mating with *Y. pseudotuberculosis* wild-type bioluminescent strain YPIII/pIBX. Selection for homologous recombination events were done on *Yersinia* agar plates, and by using sucrose pressure the *sacB* gene located on pDM4 enforced the integrated suicide plasmid to recombine out together with the chromosomal DNA to be deleted. Mutants were finally confirmed by sequencing(5).

**Bacterial RNA enrichment for total RNA extracted from *ΔssrS* infected tissue and lumen samples, RNA-seq library prepration and depletion of bacterial rRNA**

Bacterial RNA was enriched by hybridization-based depletion of host RNA, adapted from Doruk et al., 2026(6). Complementary DNA was synthesized from total RNA isolated from mesenteric lymph nodes of uninfected mice using the SuperScript IV First-Strand Synthesis System (Thermo Fisher Scientific) according to the manufacturer’s instructions. Briefly, 3 µg RNA was combined with 1 µL random hexamers (0.2 µg µL⁻¹) and 1 µL 10 mM dNTP mix in a final volume of 14 µL, incubated at 65 °C for 5 min and cooled to 4 °C for 1 min. Then 4 µL 5× first-strand buffer, 1 µL 0.1 M DTT and 1 µL SuperScript IV reverse transcriptase (200 U µL⁻¹) were added to a final volume of 20 µL. Reverse transcription was performed at 50 °C for 30 min, followed by inactivation at 80 °C for 10 min. Residual RNA template was removed by adding 2 µL RNase H (5 U µL⁻¹), 2 µL RNase A (10 mg mL⁻¹) and 2.4 µL 10x RNase H reaction buffer and incubating at 37 °C for 30 min before cooling to 4 °C. cDNA was purified with 1.8x AMPure XP beads (Beckman Coulter) to remove residual random hexamers and short cDNA fragments. Purified cDNA was used as the hybridization bait for host RNA depletion.

For depletion of host RNA, each reaction contained 300 ng total RNA from infected sampels, 750 ng cDNA, 1× hybridization buffer (50 mM Tris-HCl pH 7.5, 100 mM NaCl) and 50 µM EDTA in a final volume of 20 µL. This mixture is then subjected to a controlled hybridization protocol and reaction performed at 55°C. Total 10μL Preheated RNase H mixture (1 µL Thermostable RNase H (5U/µL, NEB M0523S), 2 µL 5x hybridization buffer, 6 μL 50 mM MgCl2, and 1 μL nuclease-free water) and then added to the sample mix and incubated at 55°C for 10 minutes. Digested samples were purified with the RNA Clean & Concentrator kit (Zymo Research) and eluted in 20 µL nuclease-free water. Residual cDNA was degraded by adding 2 µL TURBO DNase (2 U µL⁻¹; Invitrogen, AM2238) and 2.4 µL 10× TURBO DNase buffer, incubating at 37 °C for 30 min and transferring to ice. Enriched RNA was purified by a second column cleanup and quantified using the Qubit RNA HS assay (Thermo Fisher Scientific).

The enriched RNA samples were taken for library preparation using NEBNext Ultra™ II Directional RNA Library Prep kit according to the manufacturer’s protocol. Briefly, priming and fragmentation was performed by incubating RNA samples 94 °C for 7 min with random primers then transferring the tube to ice. First and second-strand cDNA synthesis, end-prep, and adaptor ligation were performed according to the manufacturer’s instructions. Finally, we constructed the libraries using the NEBNext Multiplex Oligos for Illumina kit, which is required for amplification of the ligated cDNA according to the manufacturer’s instruction. Indexed cDNA was purified 0.9x AMPure XP magnetic beads (Beckman Coulter). Libraries were quantified on a Qubit™ 4.0 fluorometer, and final library size was assessed by running libraries on Bioanalyzer High Sensitivity Chip (Agilent, 5067– 4626). As host-derived cDNA bait provides no complementarity to bacterial transcripts, bacterial rRNA was retained following host depletion. Libraries were therefore treated with the CRISPRclean Pan-Bacterial rRNA Depletion Kit (Jumpcode Genomics, San Diego, CA, USA) according to the manufacturer’s instructions to deplete bacterial rRNA-derived fragments and increase the proportion of informative mRNA reads.

**Bacterial total RNA and DNA isolation from *in vitro* samples**

Total RNA of *Y. pseudotuberculosis* was extracted by using TRIzol^TM^ (Invitrogen, Lifescience Technologies, CA, USA), according to manufacturer’s protocol. After *in vitro* treatments, cells were first centrifuged at 5000x*g* for 5 minutes, then cell pellets were resuspended in 1 mL of TRIzol, and the protocol was followed. After TRIzol extraction, upper aqueous phase was used purify total RNA via Direct-zol RNA extraction kit (Zymo Research) according to manufacturer’s protocols. To remove DNA contamination in total RNA, purified RNAs were treated with Turbo DNAse (Invitrogen, Lifescience Technologies, CA, USA) according to manufacturer’s protocols. After DNase treatment, total RNA was cleaned and concentrated via RNA clean & concentrator^TM^ kit (Zymo Research). Similarly total DNA of *Y. pseudotuberculosis* was extracted by using same TRIzol^TM^ (Invitrogen, Lifescience Technologies, CA, USA) reagent via DNA extraction protocol. After TRIzol extraction as similar with RNA extraction, lower organic phase and interphase were used to purify DNA instead of upper aqueous phase, which contains RNA, according to manufacturer’s protocol.

**cDNA Synthesis and qPCR**

Expression of 6S RNA, *yopE*, *yopD* and *yopH* genes are analysed in samples from different oxygen concentrations (0%, 0.1%, 1%, 5% and 10%) combination with T3SS induction and non-induction condition via qPCR. For each condition, DNase treated RNAs were used as a template for cDNA synthesis. For cDNA synthesis, RevertAid H minus First Strand cDNA synthesis kit (Thermo Scientific) was used. The qPCR reactions were performed in triplicate for each condition using qPCRBIO SyGreen Mix Fluorescein kit (PCR Biosystems, England) and Bio-Rad CFX Duet qPCR system. *rpoB* gene was used as internal control to calculate the relative expression of tested genes. For in vivo gene expression analysis in infected cecal tissues, total RNA isolated from WT- and ssrS-infected cecal biopsies was treated with TURBO DNase and reverse-transcribed using the RevertAid H Minus First Strand cDNA Synthesis Kit (Thermo Scientific). Bacterial *yopE*, *yopD*, and *yopH* transcript levels were quantified by qPCR and normalized to bacterial *rpoB*, while murine *Tnf*, *Cxcl2*, and *Il1b* transcript levels were normalized to host *Gapdh*. Primer sequences are listed in Supplementary Table 6.

**Western blot detection of secretion of Yop effectors**

For detection of Yops, the T3SS was induced according to previously described in different oxygen concentration. After 3 hours of induction, optical density (OD_600_) of the cultures were measured, the cells suspensions were centrifuged, and cell pellets and supernatants were separated. Supernatants were used to extract secreted proteins via precipitation with 10% TCA. Cell pellets were boiled in Laemmli sample buffer to extract cellular proteins. Extracted cellular and secreted proteins were loaded and separated in 12% SDS-PAGE, with normalization of OD_600_ values of the treated cultures. Separated proteins were transferred to Immobilon-P PVDF membrane (Millipore) and probed overnight at 4°C with a primary rabbit anti-Yops antibodies at a 1:2000 dilution. The final blot was developed by chemiluminescence (Immobilon Western Chemiluminescent HRP Substrate, Millipore) and the bands were imaged via Amersham Imager 680. Band intensities were quantified using ImageJ by measuring the signal within predefined regions of interest (ROIs).

**Determination of plasmid copy number and *in vivo* replication**

The plasmid copy number and replication rates of tissue bacteria were detected using droplet digital PCR (ddPCR) method. Plasmid copy number was determined as the number of virulence plasmid per chromosome; tissue bacterial replication rates were quantified as peak-to-through (PTR) ratio using a previously established protocol(7, 8). For this, ddPCR was performed on the DNA extracted directly from the same biopsy homogenates used for RNA extractions.

### **Lipocalin-2 quantification by ELISA**

### Cecal luminal contents from WT- and *ssrS*-infected mice at 1, 2, and 3 dpi were collected, weighed, and homogenized in sterile PBS. Homogenates were centrifuged to remove debris, and Lipocalin-2 concentrations in the supernatants were quantified using the Mouse Lipocalin-2/NGAL DuoSet ELISA kit (R&D Systems) according to the manufacturer's instructions.

### **Histology**

### For tissue preparation, mouse cecal tissues from infected mice were dissected out and immediately placed in 4% PFA for 6-12 h fixation. Tissues were rinsed with PBS and 70% ethanol followed by dehydration in Tissue-Tek^®^VIP (SAKURA). Paraffin embedding of tissues in Histowax^TM^ paraffin (Histolab) were performed with Tissue-Tek^®^TEC (SAKURA) and kept at 4 °C until sectioning. For hematoxylin and eosin staining of cecal tissues, 5 µm tissue sections were de-paraffinized by heating to 60^o^C for 20 min and 3 x 10 min xylene treatments at RT. After dehydration with decreasing concentrations of ethanol, sections were stained with Mayer hematoxylin (Histolab) and 0,2% Eosin (Histolab), and after dehydration in 99% ethanol and xylene mounted with dibutylphtalate (DPX) (Sigma Aldrich).

**Spatial gene expression**

For in-situ detection of Y. pseudotuberculosis expression of specific transcripts in foci of cecum lymphoid compartments, RNAscope staining was performed on paraffin-embedded tissue sections. The procedure employed was according to the manufacturer’s protocol (RNAscope® Multiplex Fluorescent Reagent Kit V2 (ACDBio, Bio-techne, Oxford, UK). All probes were designed by and purchased from ACDBio. To ensure probe specificity for detecting single mRNAs in Y. pseudotuberculosis, probe sequences were designed to avoid potential cross-reactivity against host cell and microbiota transcripts. For detection of the infecting bacteria a probe recognizing unique regions of Y. pseudotuberculosis YPIII 23S RNA encoded by 7 open reading frames (1 probe pair) was used. For detection of mRNA products of the virulence plasmid a probe recognizing parE (pYV0026) mRNA (1 probe pair) was used, this gene product was selected due its expression pattern in vitro with very low expression in non-inducing conditions. A probe recognizing 6S RNA was designed with 4 probe pairs and for detection of replicating bacteria a probe recognizing fis mRNA (6 probe pairs) was employed. For RNAscope labelling, 5 µm tissue sections were de-paraffinized followed by treatment with Hydrogen peroxide at RT for 10 min, Target retrieval for 10 min at 99 oC, and with Protease Plus for 15 min at 40oC in an HybEZ™ II Oven (Bio-techne). For targeting of bacterial transcripts, the specimens were incubated for 2 h with the different probes diluted in probe dilution buffer (1:100 for 23S RNA, 1:50 for 6S RNA and 1:100 for pYV0026) at 40°C in HybEZ™ II Oven. This was followed by subsequent steps for signal amplification and for fluorescent labelling of the probes, the specimens were incubated with the fluorescent dyes TSA Vivid Fluorophore 520 (32327, Bio-techne) and Opal 570 (Akoya Biosciences), both diluted 1:1500 in TSA buffer. For co-detection of Y. pseudotuberculosis transcripts and polymorphonuclear leukocytes (PMNs) in infected cecal tissues, the RNAscope protocol was supplemented with immunofluorescent staining. After final incubation of RNAscope treated specimens a protocol for immunofluorescence staining was implemented. The tissue sections were blocked subsequently by incubating in 0.1 M glycin, avidin, biotin (Vector Laboratories) and 1% BSA in PBS. Tissue sections were then stained using anti-Ly6G recognizing PMNs (clone 1A8; BD Biosciences) and a biotinylated mouse anti-rat immunoglobulin as secondary antibody (Jackson ImmunoResearch), followed by streptavidin-phycoerythrin conjugated Alexa 555 (Invitrogen). Nuclei were counterstained with 4=,6-diamidino-2-phenylindole (DAPI) provided in the RNAScope kit. The specimen were mounted with ProLong Gold Antifade Mountant (Invitrogen). Images were captured using a Hamamatsu Orcha C4742-95 camera and NIS-Elements AR (version 3.2) software (Nikon Instruments).

**Microscopy image analysis**

The microscopy image analysis was divided into three different parts: colocalization (as in Supplementary Fig. S11a), area (as in Fig. 6d and Supplementary Fig. S11b), and line plot/distance analysis (as in Fig. 6e and 6f). As the microscope magnification varied between 10x and 20x between different samples, all samples were scaled to a 20x reference pixel frame. The colocalization analysis was performed using a custom ImageJ macro that automated the processing of all composite images First, channels were split and background subtraction was applied to both red and green channels using the rolling ball algorithm (radius = 50 pixels). Otsu thresholding was then applied individually to each channel to create binary masks encompassing significant fluorescence signals. The masks were combined using a logical OR operation to define regions of interests (ROIs) containing bacterial populations with segmentation, with analysis restricted to objects above 5000 pixels to exclude noise. Within these ROIs, mean thresholding was applied to both channels to establish optimal signal detection thresholds for colocalization analysis. The colocalization metric “Pearson’s correlation coefficient” was calculated using the Coloc2 plugin in ImageJ. Statistical comparisons of Pearson correlation coefficients were made across different RNA target conditions to assess colocalization patterns between RNA targets and 23S RNA control signals.

### The area analysis was performed using another custom ImageJ macro to quantify the spatial heterogeneity of each target RNA. For each image, channels were automatically split and processed using Otsu thresholding to generate binary masks for red (RNA target) and green (23S rRNA - control) bacterial signals. Afterwards the overlap of both masks was quantified using the intersection of red and green masks. This overlap was then divided once by the masked green area and once by the masked red area. The ratio “intersection / green”, gives an indication on how homogenous the target RNA is expressed in the bacterial control group. A high number of this ratio signals a very homogenous distribution. The ratio “intersection / red” serves as a control, that the RNA target is mainly co-expressed with the control RNA. For statistical analysis, all results from one RNA target are batched and visualised in a box plot to compare between the different target RNAs.

### For the line/distance analysis, for each sample 3-5 intensity plots were extracted across all three channels (DAPI, 23S rRNA and target) in ImageJ by manually drawing lines orthogonal to the tissue/lumen border. Afterwards each intensity profile was smoothed by a gaussian filter and normalized with channel-wise min-max normalization. The red channel (target channel) was normalized to the min/max of the green channel to ensure comparisons between different RNA targets. The border was determined and saved by manually detecting the location of the tissue/lumen border position within the line plot. This was assessed by eye at the position, where the DAPI signal abruptly increased. As a final step, the ratio between the smoothed and normalized RNA target and control was calculated. This analysis (acquisition of line profile, smoothing, normalizing and ratio calculation) was done for every line (n=150) of every sample (n=43) and then averaged by sample and RNA target.

### **Single-colony growth analysis by HeteroTyper imaging**

At 2 dpi, WT and *ΔssrS* from tissue and lumen were dissected and homogenized in phosphate-buffered saline (PBS). Homogenates were serially diluted in PBS and 100 µL of dilutions (from 10^1^-10^3^) were spread onto LA plates. The plates were incubated at room temperature (RT) for 20 hours (h) and imaged in the HeteroTyper system to monitor colony growth for 28h with 30 minutes (min) intervals(9).

For *in vitro* growth experiments, the WT and *ΔssrS* strains were subcultured in LA supplemented with kanamycin (50 µg/mL). The strains were cultured overnight at 26°C and 160 rpm in LB. The overnight culture was co-cultured in the morning to a starting optical density at 600 nm (OD_600_) of 0.05 and grown in LB at 26°C and 160 rpm. Samples were collected at different time points (24, 48, and 72 hours) during growth. Collected cultures were then serially diluted, and 100 µL of dilutions (from 10^4^-10^6^) were spread onto 6 LA plates for each strain and growth period. The plates were incubated at RT for 20h and imaged in the HeteroTyper system for 40h with 30min intervals. For *in vitro* stress experiments, the WT and *ΔssrS* strains were grown following the same protocol as the *in vitro* growth experiments but collected at 5 hours of growth. The cultures were exposed to different *in vivo* relevant stress conditions, namely acidic stress (10% acetic acid v/v), low iron (1 mM 2,2’-Bipyridyl; Sigma-Aldrich), osmotic stress (0.5 M NaCl), and oxidative stress (1 mM H_2_O_2_), including an unexposed control condition. The cultures were incubated at 26°C and 160 rpm. Following 10 minutes stress exposure, samples were collected, serially diluted and spread onto 6 LA plates for each condition. The plates were incubated at RT for 16h and imaged in the HeteroTyper system for 29h with 30min intervals.

**Image processing and colony features**

To compare bacterial growth dynamics across experimental setups, agar plate images acquired with the HeteroTyper system were processed with the integrated downstream pipeline(9). Each experimental setup was analysed individually. Colony appearance time (h) was quantified from the smoothed time-course colony areas generated by the pipeline. Maximum growth rate (pixels per hour, px/h) was estimated for each colony by fitting a Gompertz model to the smoothed colony area over time. Colony appearance times were visualised as ridgeline plots and the relationship between colony appearance time and maximum growth rate was visualised using scatter plots. The analyses were performed in MATLAB R2026a.

**Supplementary Figures**
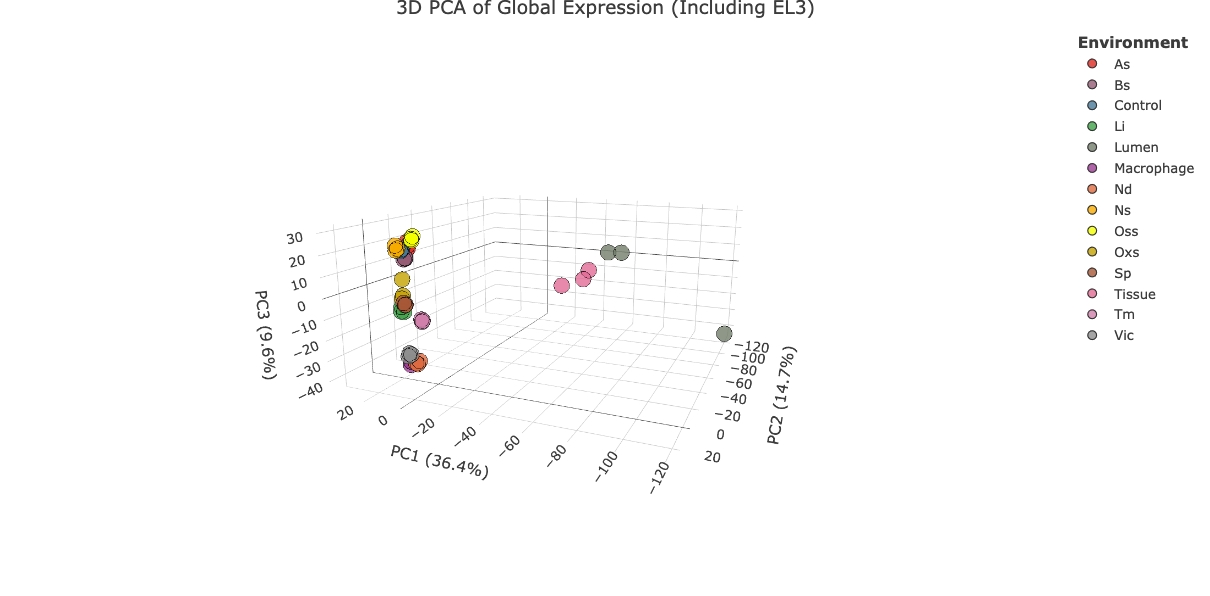

**Supplementary Figure S1. Principal component analysis (PCA) of Yersinia pseudotuberculosis transcriptomes under infection-relevant stress conditions and *in vivo* environments** **with distant clustering of *in vivo* transcriptomes**. PCA was performed using TPM values from the PATHOgenex dataset (Avican et al., 2021), which includes RNA-seq data from bacteria exposed to various *in vitro* stressors along with control condition. Each *in vitro* condition includes three biological replicates. Additionally, transcriptomes from bacteria isolated from cecal lumen and tissue of three infected mice are shown. Different stressors are acidic stress (As), bile stress (Bs), control (Ctrl), low iron (Li), microaerophilic growth/hypoxia (Mig), nutritional downshift (Nd), nitrosative stress (Ns), osmotic stress (Oss), oxidative stress (Oxs), stationary phase (Sp), temperature (Tm), and virulence-inducing condition (Vic).

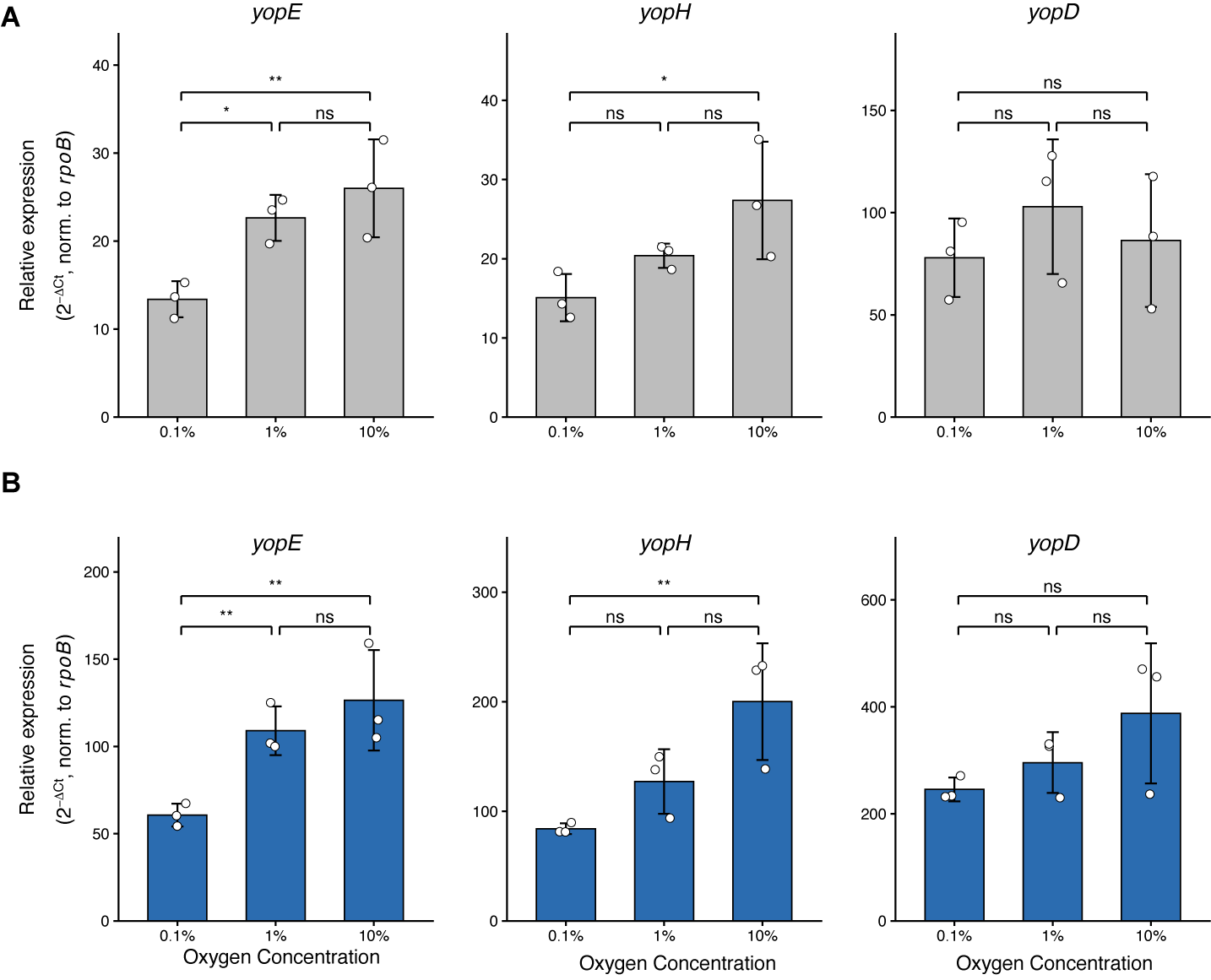

**Supplementary Figure S2. The expression of T3SS genes increases upon increased oxygen concentartion. A-B** Relative expression of *yopE*, *yopH*, and *yopD* under T3SS non-inducing (37°C) **(A)** and inducing (37°C, Ca^2+^ depletion) condition **(B)** in 0,1%, 1%, and 10% oxygen concentrations for 3 hours measured with qPCR. Relative expression of *yopE*, *yopH*, and *yopD* are calculated according to Ct values normalized to that of *rpoB*. All qPCR experiments are performed with three biological and three technical replicates. Data are presented as mean values ± SD from 3 sets (n=3) for each condition. Statistical significance was assessed by one-way ANOVA comparing oxygen concentrations within each condition, inducing and non-inducing analysed separately. Asterisks represent p-value thresholds: **** for p < 0.0001, *** for p < 0.001, ** for p < 0.01, and * for p < 0.05; ns, non significant.

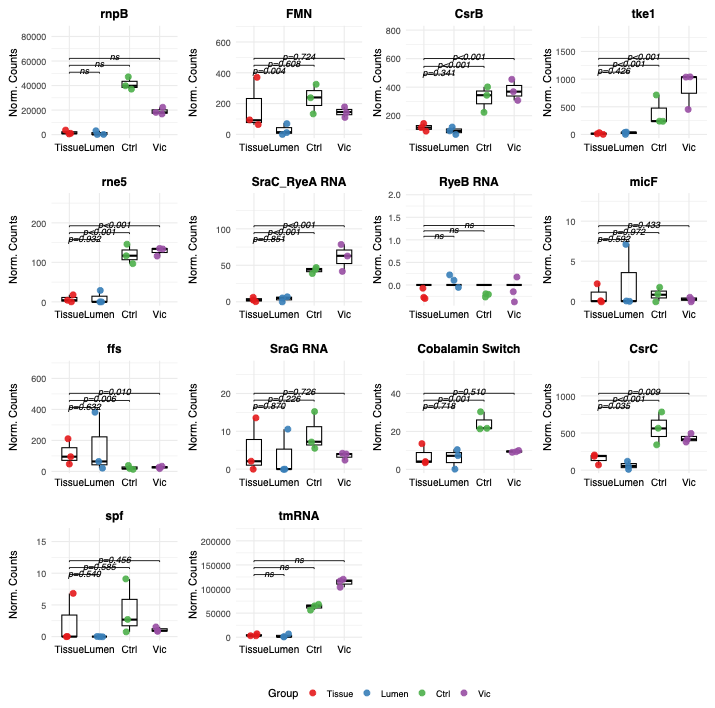

**Supplementary Figure S3. Expression levels of targeted ncRNAs across *in vivo* and *in vitro* environments.**Expression levels of 14 ncRNAs were assessed in lumen- and tissue-located bacteria, as well as in *in vitro* environmental (26°C, Ctrl) and virulence-inducing (T3SS, Vic) conditions from the PATHOgenex dataset (Avican et al., 2021). Gene expression is presented as DESeq2-normalized counts to account for library size and biological variance. Data are visualized as boxplots (center line, median; box limits, upper and lower quartiles) overlaid with individual data points representing independent biological replicates (n=3 per condition). Statistical significance was calculated using a negative binomial generalized linear model with empirical Bayes variance shrinkage (native DESeq2 Wald test). P-values displayed above the brackets indicate pairwise comparisons of the host tissue environment against the lumen, Ctrl, and Vic conditions.

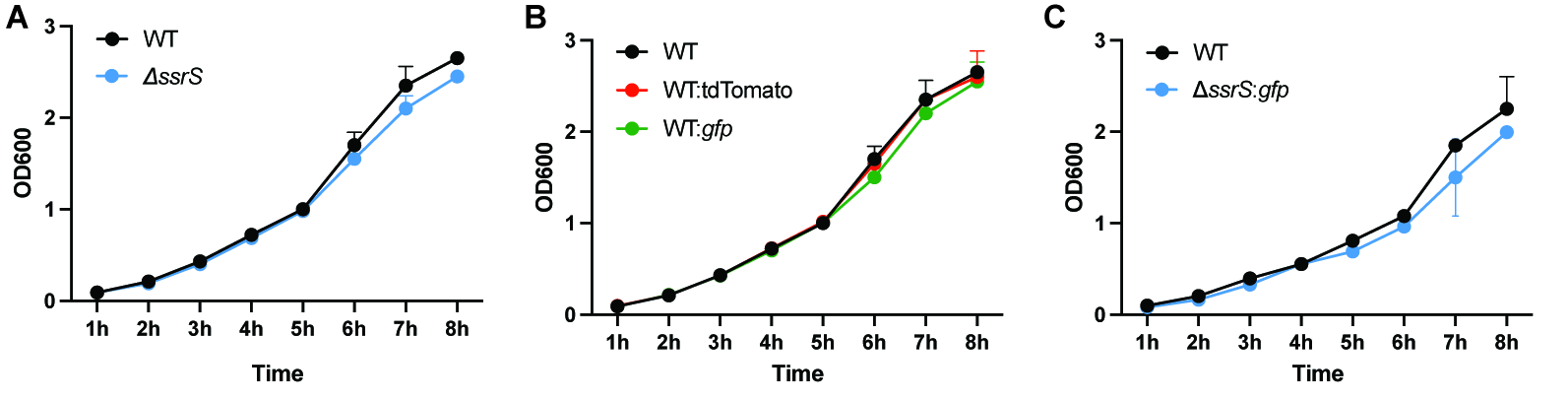
**Supplementary Figure S4.** Growth curves of Y. pseudotuberculosis wild-type and mutant strains used in this study. All strains exhibited comparable growth dynamics, reaching stationary phase with no or negligible differences (slight growth defect for *ssrS* mutant) in growth rates, indicating that the introduced mutations and reporter constructs did not affect general fitness under these conditions. Cultures were initiated with 1:100 dilutions of overnight cultures and grown in LB medium at 26 °C with aeration for 8 hours. Optical density at 600 nm (OD₆₀₀) was measured hourly to monitor bacterial growth. **A** WT and *ssrS* deletion mutant. **B** WT, WT:*tdTomato*, and WT:*gfp*. **C** WT and *ssrS:gfp*. Three biological replicates (n=3) were used in **A**, **B**, and **C**.

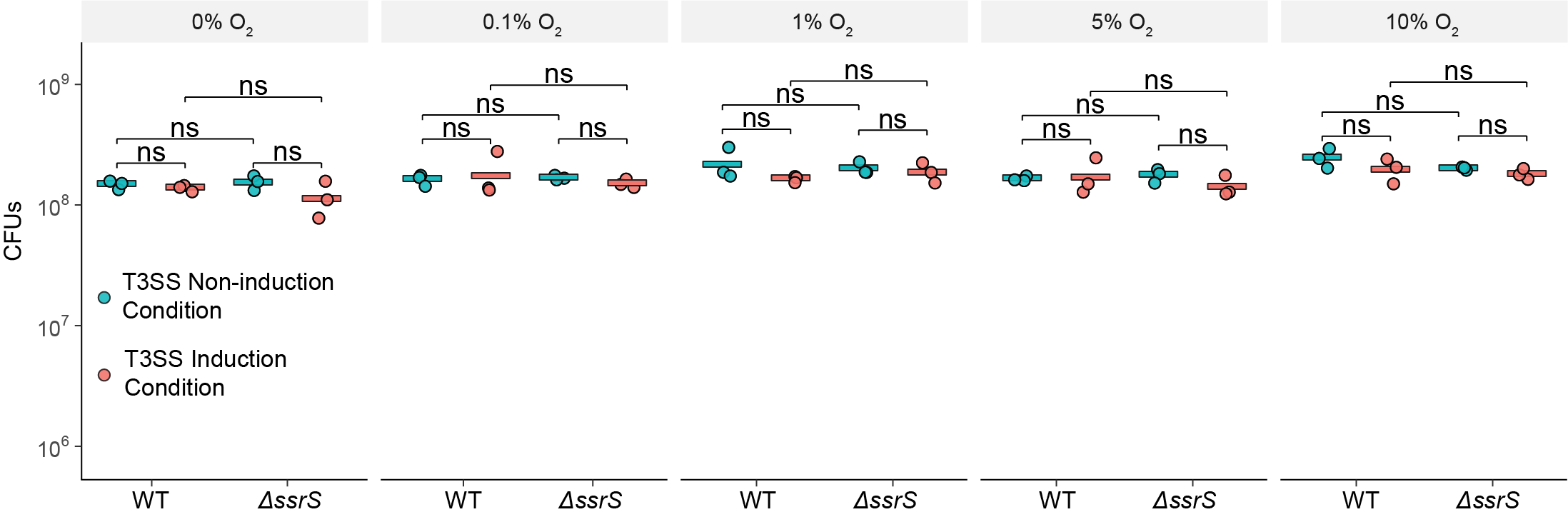

**Supplementary Figure S5: Loss of 6S RNA (ΔssrS) does not affect viability of Yersinia pseudotuberculosis across a range of oxygen tensions under T3SS-inducing and non-inducing conditions.** Wild-type (WT) and 6S RNA-deficient (ΔssrS) strains were grown at the indicated oxygen concentrations (0%, 0.1%, 1%, 5%, and 10% O₂) under T3SS non-induction (teal) and T3SS induction (salmon) conditions, and viability was determined as colony-forming units (CFUs). Each point represents an independent biological replicate (n = 3) and the horizontal bar indicates the geometric mean. The y-axis is shown on a log₁₀ scale. Statistical significance between the indicated groups was assessed by pairwise two-sample t-tests on log₁₀-transformed CFU values-, ns, not significant (P > 0.05).

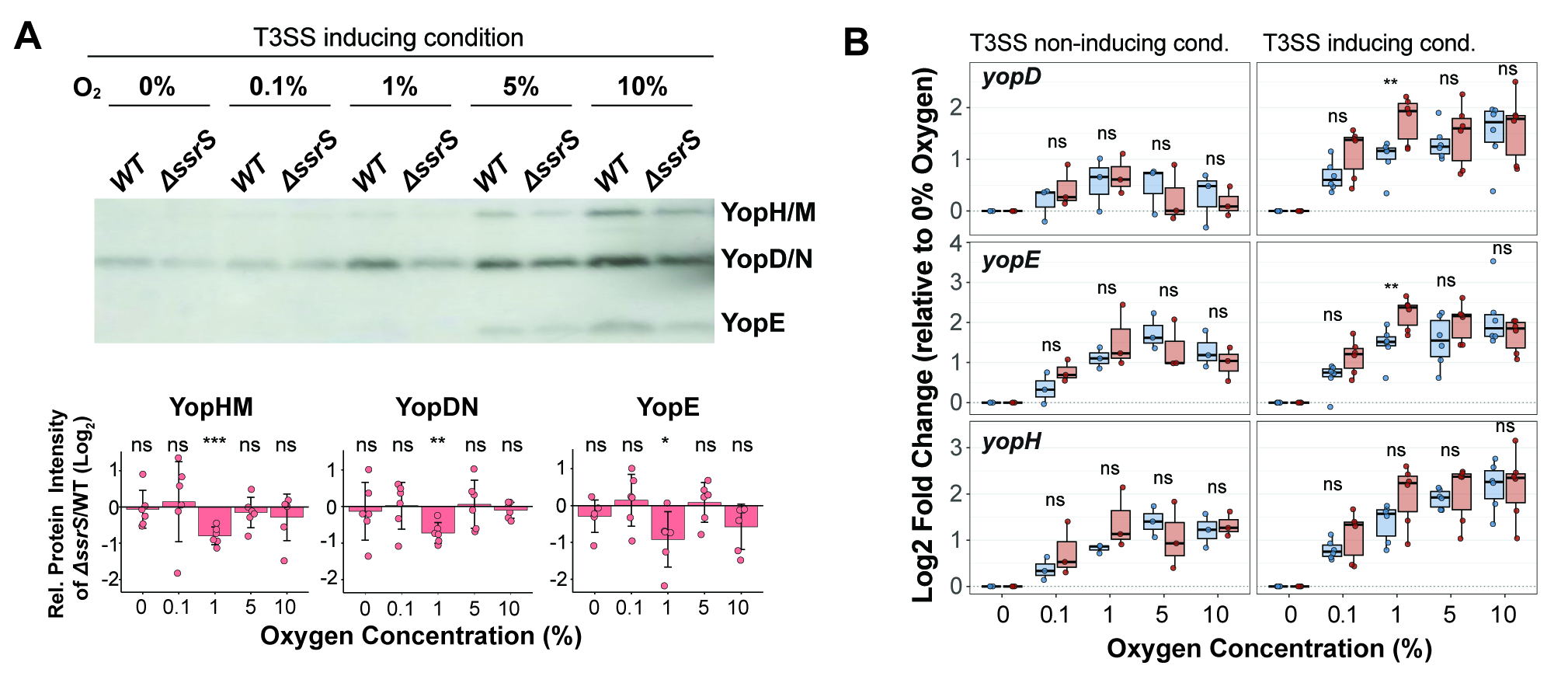

**Supplementary Figure S6. T3SS in the loss of 6S RNA. A** Secretion of *Y. pseudotuberculosis* T3SS effectors under T3SS inducing in different oxygen concentrations for 3 hours (upper panel). Amounts of proteins loaded were normalized according to CFUs counted after 3 hours. Log₂‑transformed relative Yop intensities (lower panel) were calculated by normalizing Δ*ssrS* Yop intensities to WT Yop intensities. Data are presented as mean values ± SD from 6 sets (n=6) for each condition. One-sample t tests were performed on log₂‑transformed relative intensity values. Asterisks represent p‑value thresholds: **** for p < 0.0001, *** for p < 0.001, ** for p < 0.01, and * for p < 0.05; ns, non significant. **B** Differential expression of *yopH, yopD,* and *yopE* in ∆*ssrS* and in WT under T3SS inducing condition in different oxygen concentrations measured with qPCR. Log2 transformed fold changes were calculated as Log_2_(2^-ΔΔCT^) values using *rpoB* as reference gene and 0% Oxygen concentration as control. All qPCR experiments are performed with three biological and three technical replicates. Data are presented as mean values ± SD from 3 sets (n=3) for each condition. One‑sample t‑tests were performed on Log_2_(2^-ΔΔCT^) values to assess differences relative to the 0% oxygen control. Asterisks represent p‑value thresholds: **** for p < 0.0001, *** for p < 0.001, ** for p < 0.01, and * for p < 0.05; ns, non significant.

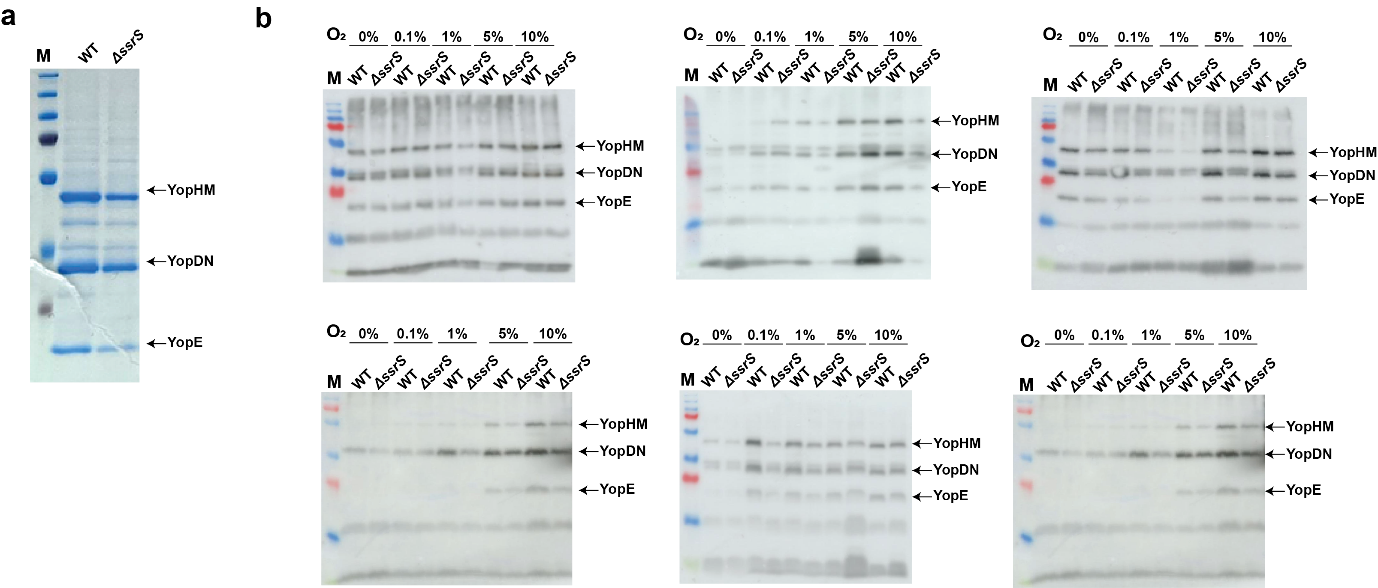

**B**

**A**

**Supplementary Figure S7. T3SS secretion in Y. pseudotuberculosis WT and *ΔssrS* mutant strains under T3SS-inducing condition at varying oxygen levels.** **A** SDS-PAGE analysis of secreted proteins (secretome) from WT and *ΔssrS* strains under T3SS-inducing conditions (37 °C, Ca²⁺ depletion) at atmospheric oxygen levels (21%) for 3 hours. **B** Western blot detection of secreted T3SS effector proteins was conducted in six independent replicates (n = 6) for WT and *ΔssrS* strains under T3SS‑inducing conditions (37 °C, Ca²⁺ depletion) after 3 hours at low oxygen concentrations (0%, 0.1%, 1%, 5%, and 10%).

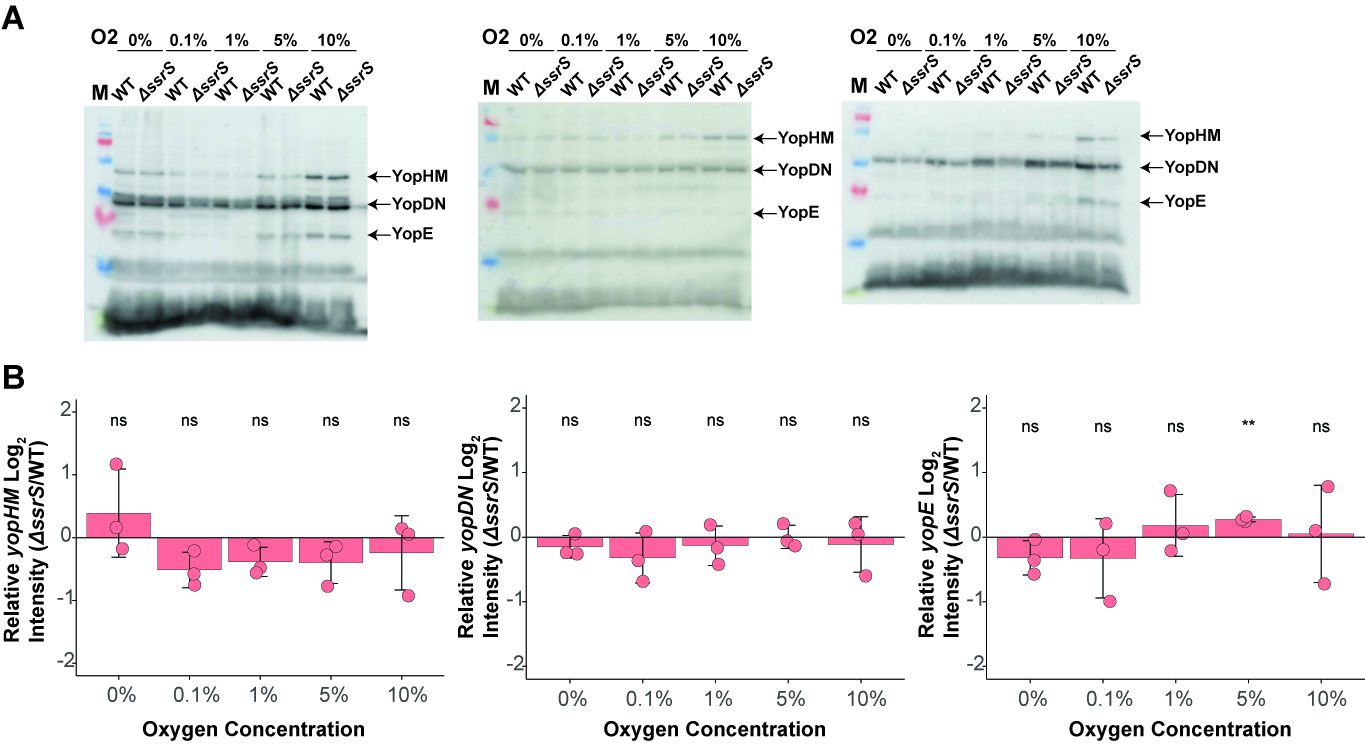

**Supplementary Figure S8. T3SS expression in Y. pseudotuberculosis WT and *ΔssrS* mutant strains under T3SS-inducing conditions at varying oxygen levels.** **A** Western blot detection of expressed T3SS effector proteins was conducted in three independent replicates (n=3) for WT and *ΔssrS* strains’ pellets under T3SS-inducing conditions (37 °C, Ca²⁺ depletion) across a range of low oxygen concentrations (0%, 0.1%, 1%, 5%, and 10%) for 3 hours. **B** Log₂‑transformed relative protein intensities were calculated by normalizing *ΔssrS* protein intensities to WT protein intensities according to three sets of western blotting (n=3). One-sample t tests were performed on log₂‑transformed relative intensity values. Asterisks represent p value thresholds: **** for p < 0.0001, *** for p < 0.001, ** for p < 0.01, and * for p < 0.05; ns, nonsignificant.

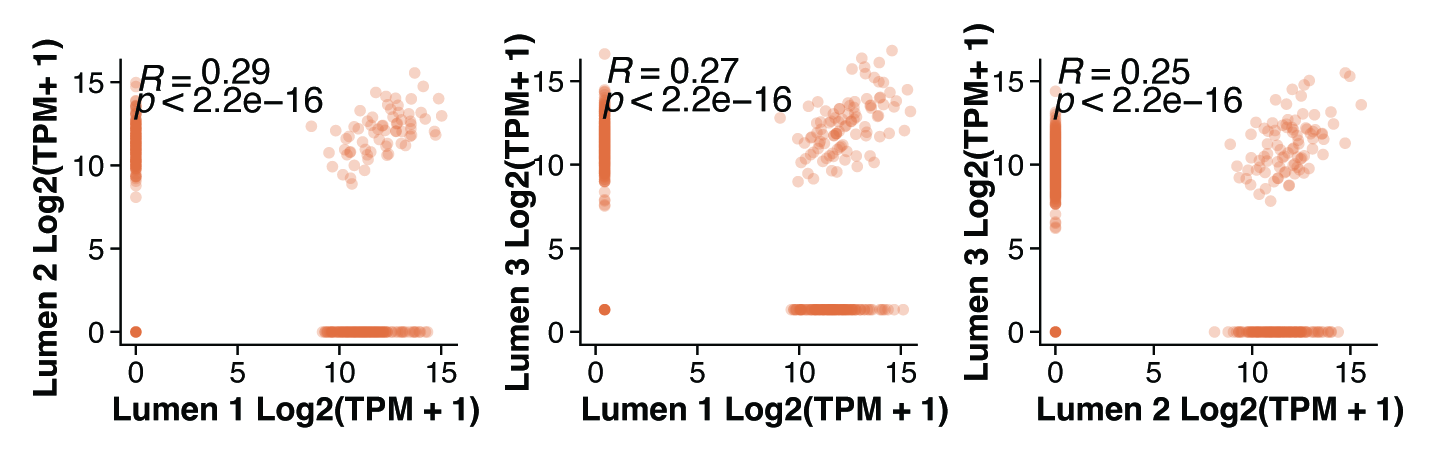

**Supplementary Fig. S9. Pairwise correlation of *ΔssrS* luminal transcriptomes.** Scatter plots of log₂(TPM + 1) values for all annotated CDSs between pairs of *ΔssrS* luminal replicates (Lumen 1 versus Lumen 2, Lumen 1 versus Lumen 3, and Lumen 2 versus Lumen 3). Each point represents one gene; correlation coefficients (R) and p values are indicated.

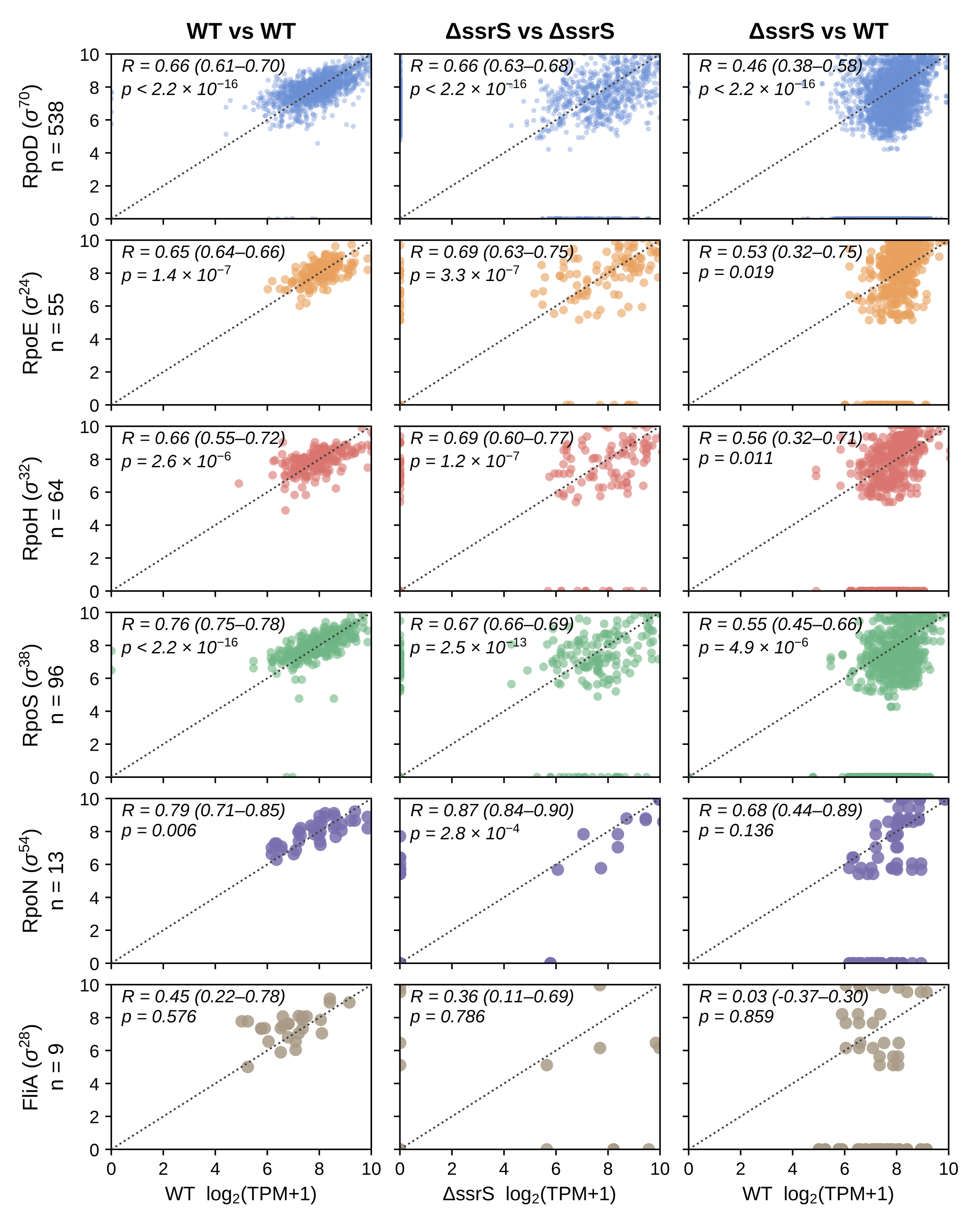

**Supplementary Figure S10. Sigma factor regulon expression in WT and *ΔssrS* tissue bacteria, within and between genotypes.** Pairwise scatter plots of gene expression for each sigma factor regulon in *Y. pseudotuberculosis* recovered from infected caecal lymphoid tissue. Each point is one gene in one replicate pair. Rows correspond to the six sigma factor regulons tested, with the number of orthologues (*n*) analysed given at left. Columns compare biological replicates of the same genotype (WT vs WT, three pairs from three replicates; *ΔssrS* vs *ΔssrS*, three pairs from three replicates) or replicates of different genotypes (*ΔssrS* vs WT, nine pairs). In the third column the y axis is *ΔssrS* and the x axis is WT. All replicate pairs are overlaid within each panel, so panels in the third column contain three times as many points as those in the first two. R is the mean Spearman correlation coefficient across replicate pairs, with the range across pairs given in parentheses. The p value shown is the largest obtained across those pairs and is therefore the most conservative of the pairwise tests. RpoN and FliA retained too few high-confidence orthologues (n = 13 and 9) for reliable estimation and are shown for completeness only.

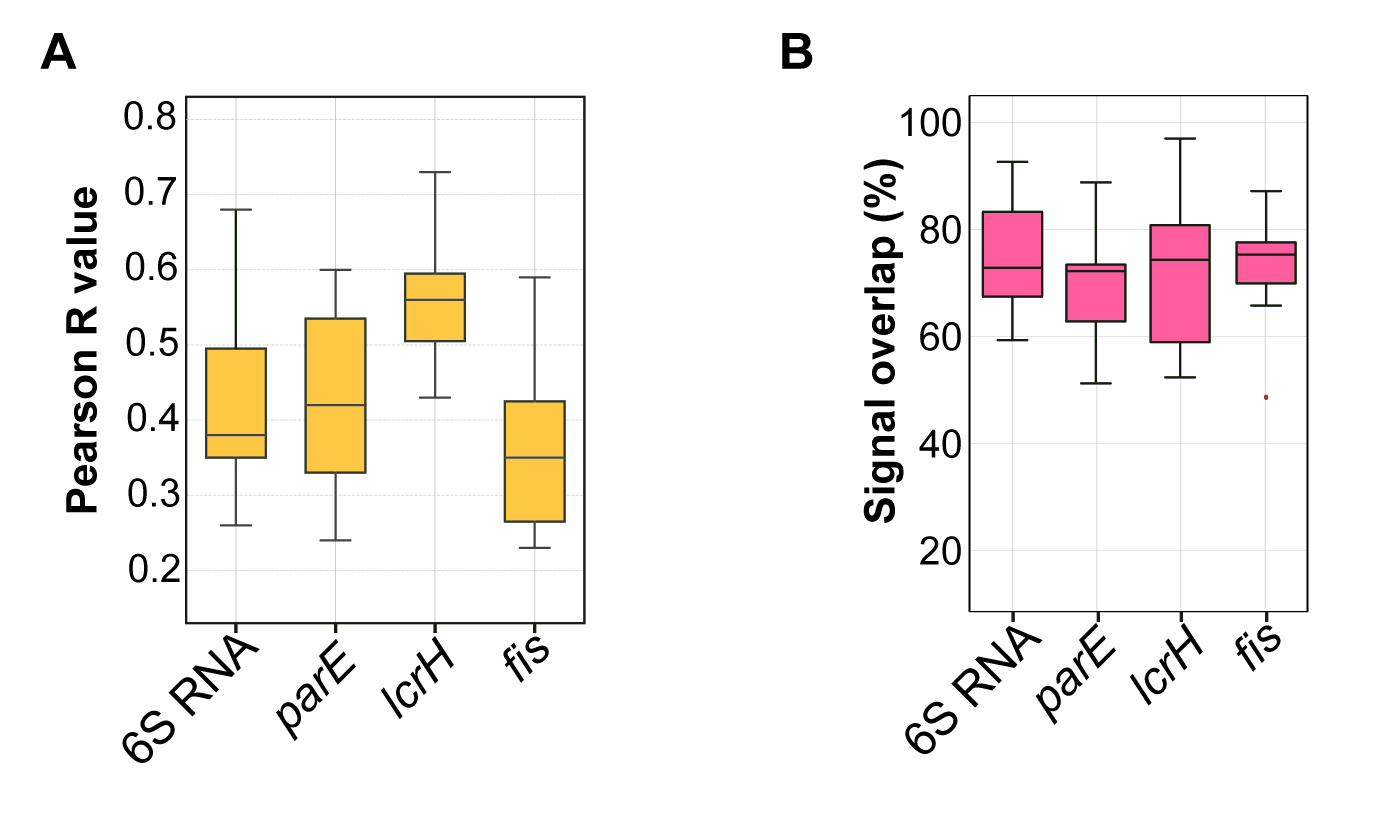

**Supplementary Figure S11. Computational analysis of RNAscope assay reveals heterogeneous expression of 6S RNA, parE, lcrH, and fis in Y. pseudotuberculosis**. **A** Pearson correlation analysis between green fluorescence signals from RNAscope probes targeting Y. pseudotuberculosis 23S rRNA and red signals from probes targeting 6S RNA, parE, lcrH, and fis. **B** Quantification of the percentage of between red signals (6S RNA, parE, lcrH, and fis) that spatially overlap with green 23S rRNA signals, indicating the degree of transcript co-localization within bacterial cells.

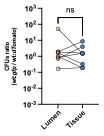

**Supplementary Figure S12. Co-infection of mice with WT *Y. pseudotuberculosis* strains ectopically expressing flurescent reporter proteins do not have an effect on infection outcome.** Ratio of CFUs recovered from cecal tissue and lumen compartments of mice (n=8) co-infected with 1:1 mixture of *Y. pseudotuberculosis* WT:*gfp* and *Y. pseudotuberculosis* WT: *tdTomato* strains. The two strains were distinguished with fluorescent signal by GFP expression and by tdTomato expression in WT strain. Statistical significance between two groups was calculated via Mann-Whitney test.

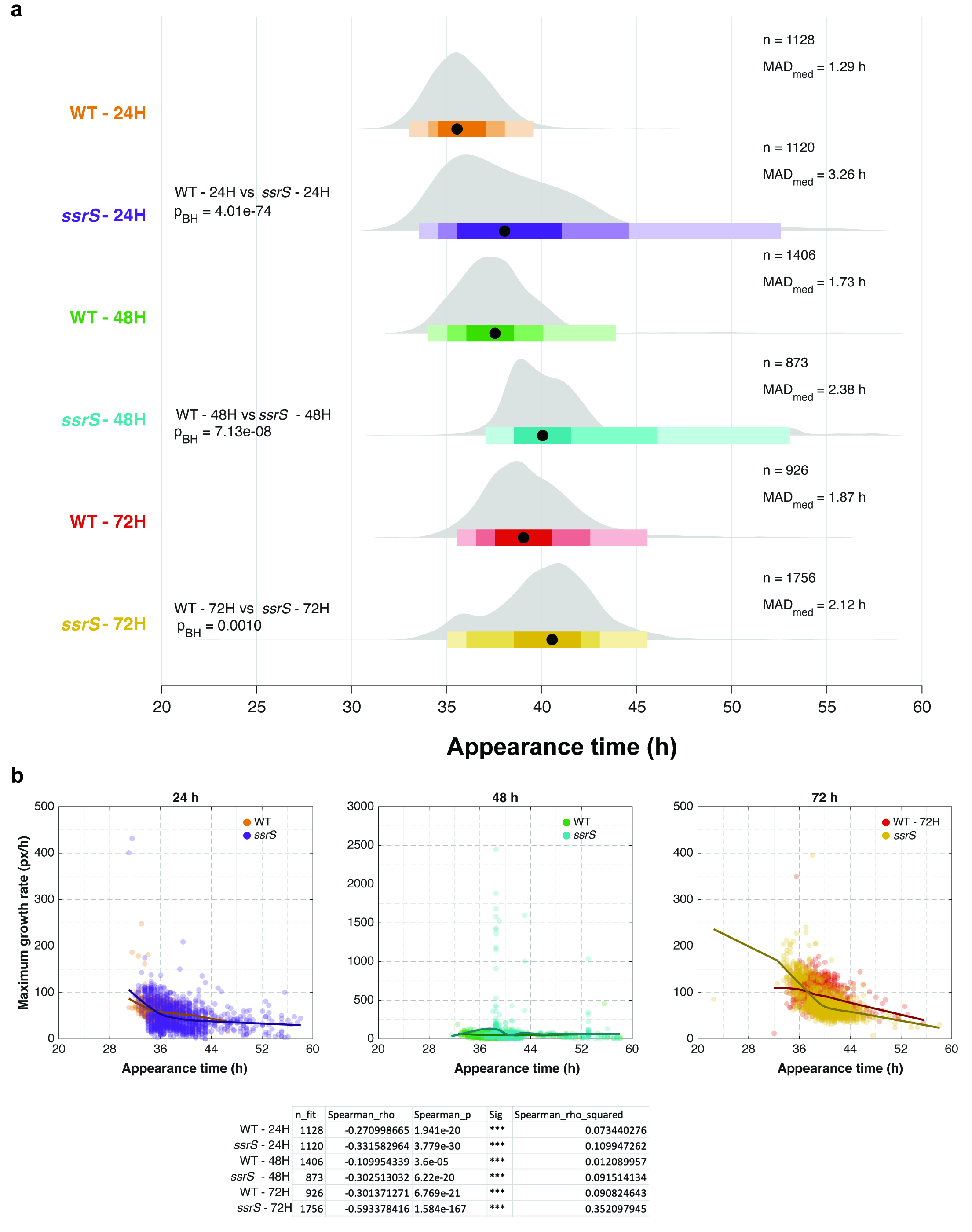

**Supplementary Figure S13. Single-colony appearance time and maximum growth rate of WT and Δ*ssrS* bacteria after 24, 48 and 72 h of culture.** WT and *ΔssrS* were inoculated into a rich liquid medium and grown at 26°C. Samples were collected from the same culture at 24, 48, and 72 hours of growth, serially diluted and spread onto 6 plates for each time point. Plates were incubated at room temperature for 20 hours and imaged in the HeteroTyper system for 40h at 30 minutes intervals. The number of colonies (n) analysed were as follows: WT, 1128 (24h), 1406 (48h), and 926 (72h); *ΔssrS*, 1120 (24h), 873 (48h), and 1756 (72h). **a** Colony appearance time shown as density ridgeline plots. Black dots below each filled area represent the group median, and bars show the central 50% (interquartile range, IQR; darkest), 80%, and 95% (lightest) of colonies. The vertical dashed line marks the median of all groups. The significance of differences in appearance time distributions between strains at each growth period was tested using the Brown-Forsythe test followed by ANOVA and corrected using Benjamini-Hochberg (BH). n, median appearance time (h), mean absolute deviation from the group median (MAD_med_), and Gini coefficient are reported per group. **b** Colony appearance time versus maximum growth rate (px/h), estimated by fitting a Gompertz model to colony area over time, for colonies of WT and *ΔssrS* at 24h (left panel), 48h (middle panel), and 72h (right panel), are represented as scatter plots. Each dot represents an individual colony. Locally estimated scatterplot smoothing (LOESS) curves are colored by respective strain and growth period. n, Spearman correlation coefficients (*ρ*), *p*-values (*p*), statistical significance asterisks, and squared Spearman correlation coefficients are represented (*ρ*^2^) are given in the table below panel **b** for each strain and growth period. Appearance time is represented in hours (h), while the colony size was measured in pixels (px).

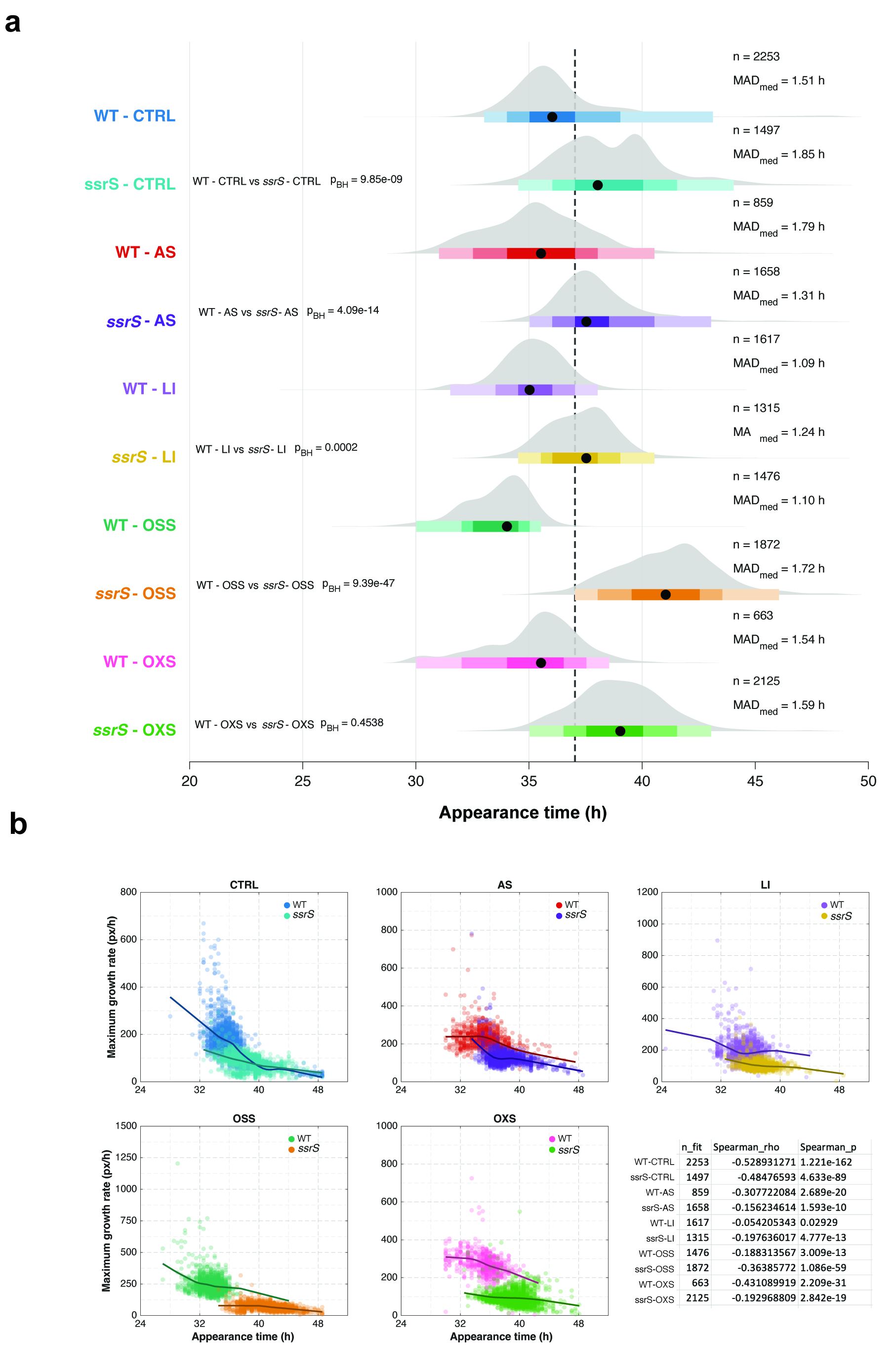

**Supplementary Figure S14. Single-colony appearance time and maximum growth rate of WT and Δ*ssrS* bacteria under acid, iron-limiting, osmotic and oxidative stress.** WT and *ΔssrS* were inoculated into rich liquid medium and grown at 26°C for 5 hours. Cells were exposed to stressors for 10 minutes and were spread onto 6 plates for cells exposed to acidic stress (AS), low iron (LI), osmotic stress (OSS), oxidative stress (OXS), and a non-exposed control group (CTRL). Samples were serially diluted and spread onto 6 plates for each time point. Plates were incubated at room temperature for 16 hours and imaged in the HeteroTyper system for 29h at 30 minutes intervals. **a** Colony appearance time shown as density ridgeline plots. Black dots below each filled area represent the group median, and bars show the central 50% (interquartile range, IQR; darkest), 80%, and 95% (lightest) of colonies. The vertical dashed line marks the median of all groups. The significance of differences in appearance time distributions between strains for each growth condition was tested using the Brown-Forsythe test followed by ANOVA and corrected using Benjamini-Hochberg (BH). Number of colonies analysed (n), median appearance time (h), mean absolute deviation from the group median (MAD_med_). **b** Colony appearance time versus maximum growth rate (px/h), estimated by fitting a Gompertz model to colony area over time, is represented as scatter plots for WT and *ΔssrS* colonies under unexposed CTRL, AS, and LI (top, left to right), and OSS and OXS (bottom, left and middle). n, Spearman correlation coefficients (*ρ*) and *p*-values (*p*) are given in the bottom-right panel in **b**. Each dot represents an individual colony. Locally estimated scatterplot smoothing (LOESS) curves are colored by respective strain and growth condition. Appearance time is represented in hours (h), while the colony size was measured in pixels (px).
